# Pyk2 in D1 receptor-expressing neurons of the nucleus accumbens modulates the acute locomotor effects of cocaine

**DOI:** 10.1101/729848

**Authors:** Benoit de Pins, Enrica Montalban, Peter Vanhoutte, Albert Giralt, Jean-Antoine Girault

## Abstract

The striatum is a critical brain region for locomotor response to cocaine. Although the D1 receptor-expressing neurons are centrally involved in mediating the locomotor effects of cocaine, the molecular pathways controlling this response are not fully understood. Here we studied the role of Pyk2, a non-receptor calcium-dependent protein-tyrosine kinase, in striatum-related functions. We discovered that cocaine injection increases Pyk2 phosphorylation in the striatum of mice *in vivo*. Pyk2-deficient mice displayed an altered locomotor response to acute cocaine injection. In contrast, they developed normal locomotor sensitization and cocaine-conditioned place preference. Accordingly, a cocaine-activated signaling pathway essential for these late responses, ERK phosphorylation, was not altered. Specific deletion of Pyk2 in the nucleus accumbens or in D1 neurons reproduced this phenotype, whereas deletion of Pyk2 in the dorsal striatum or in A2A receptor-expressing neurons did not. Mice lacking Pyk2 in D1-neurons also displayed lower locomotor response to the D1 receptor agonist SKF-81297 but not to an anticholinergic drug. Our results identify Pyk2 as a regulator of acute locomotor responses to psychostimulants and suggest that changes in Pyk2 expression or activation may alter specific responses to drugs of abuse, or possibly other behavioral responses linked to dopamine action.

## Introduction

The striatum is the main input structure of the basal ganglia, involved in motor coordination, action selection, and motivation. The GABAergic medium-size, spiny striatal projection neurons (SPNs) constitute the vast majority of striatal neurons (∼95% in rodents). They integrate cortical and thalamic excitatory inputs and are modulated by dopaminergic inputs and striatal interneurons [1]. Two major populations of SPNs can be distinguished according to their projection targets and their molecular profile [1–3]. SPNs enriched in enkephalin, D2 dopamine (D2R), and A2A adenosine (A2AR) receptors project to the external globus pallidus (GPe) thus participating in the indirect striatonigral pathway, while SPNs enriched in substance P, dynorphin, and D1 dopamine receptor (D1R) innervate the internal globus pallidus (GPi) and the substantia nigra pars reticulata (SNr), forming the direct pathway [1,2,4]. Striatal dysfunction is associated with several pathologies ranging from movement disorders in Parkinson’s or Huntington’s diseases to addiction [5]. The striatum is a major site of action of drugs of abuse which all share the ability to increase extracellular dopamine levels [6]. The signaling pathways involved in the action of glutamate, dopamine, and other neurotransmitters in SPNs have been extensively studied [7]. Although attention mostly focused on pathways involving protein serine/threonine phosphorylation, tyrosine phosphorylation is also likely to be important [8]. The regulation and functional importance of tyrosine kinases in striatal neurons are still poorly characterized.

Pyk2 is a Ca^2+^-dependent non-receptor tyrosine kinase highly expressed in forebrain neurons [9] where it is activated by neuronal activity and excitatory neurotransmission [10–12]. Pyk2 can associate with the NMDA receptor complex and has a role in synaptic plasticity [11,13–15]. Ca^2+^ triggers Pyk2 autophosphorylation on Tyr-402, which recruits and activates Src-family kinases (SFKs) [16,17]. In turn, SFKs phosphorylate other residues in Pyk2 and associated proteins, and initiate multiple signaling pathways. The striatal-enriched protein tyrosine phosphatase (STEP) dephosphorylates Pyk2 [18]. Activated Pyk2 regulates many cellular functions [19,20] and Pyk2 is associated with several pathologies including cancer [21], inflammatory diseases [20], Huntington’s [15] and Alzheimer’s diseases [22–25]. Work with Pyk2 knockout mice showed that Pyk2 in the hippocampus is involved in spatial memory and regulates spine density and morphology, as well as long-term potentiation [15] and depression [26]. These studies illustrate the role of Pyk2 in synaptic functions and behavior in physiological and pathological conditions.

Here, we use constitutive and conditional KO mice to explore Pyk2 function in the striatum. We find that Pyk2 is preferentially expressed in D1 receptor-expressing SPNs in the ventral striatum. Pyk2 phosphorylation is increased following cocaine injection. Deletion of Pyk2 reduces the acute locomotor response to cocaine but, surprisingly, neither locomotor sensitization nor conditioned place preference. Use of conditional deletion shows a specific involvement of Pyk2 in D1 receptor-expressing SPNs of the nucleus accumbens (NAc) in this response.

## Materials and Methods

### Animals

Floxed Pyk2 mice (Pyk2^f/f^) were generated by the insertion of LoxP sequences (Gen-O-way, Lyon, France) surrounding *PTK2B* exons 15b-18 coding for the kinase domain [27]. Homologous recombination was carried out in C57/Bl6 embryonic stem cells and germline transmission of the mutated allele was achieved in the same background. Floxed Pyk2 mice were initially bred to Cre-deleter mice to generate constitutive knockout mice, or to *Drd1::Cre* mice, Tg(Drd1a-cre)EY262Gsat [28], or *Adora2a::Cre* mice, Tg(Adora2a-cre)2MDkde [29], to generate conditional KO mice (Pyk2^f/f;D1::Cre^ and Pyk2^f/f;A2A::Cre^, respectively). Mice were housed at 19–22 °C with 40–60% humidity, under a 12:12 h light/dark cycle, and had ad libitum access to food and water. Animal experiments and handling were in accordance with ethical guidelines of Declaration of Helsinki and NIH, (1985-revised publication no. 85-23, European Community Guidelines), and French Agriculture and Forestry Ministry guidelines for handling animals (decree 87849, licence A 75-05-22) and approval of the Charles Darwin ethical committee project #2016111620082809. Mice used in this study were 3-6-month-old males.

### Viral vectors and stereotaxic injection

For deletion of Pyk2 in the NAc or dorsal striatum (DS), 3-month Pyk2^f/f^ mice were stereotaxically injected with AAV expressing Cre recombinase (AV-9-PV2521, AAV9.CamKII.HI.eGFP-Cre.WPRE.SV40, Perelman School of Medicine, University of Pennsylvania, USA), referred to below as AAV-Cre. As control, we injected AAVs expressing GFP (AV-9-PV1917, AAV9.CamKII0.4.eGFP.WPRE.rBG, same source), referred to below as AAV-GFP. Following anesthesia with pentobarbital (30 mg kg^−1^), we performed bilateral stereotaxic injections of AAV-GFP, AAV-GFP-Cre or AAV-GFP (2.6 × 10^9^ GS per injection) in the NAc at the following coordinates from the bregma (millimeters), anteroposterior, 1.3, lateral, ±1.3, and dorsoventral, −4.5 or in the DS, anteroposterior, 0.9, lateral, ±1.5, and dorsoventral, −2.75. AAV injection was carried out in 2 min. The cannula was left in place for 5 min for complete virus diffusion before being slowly pulled out of the tissue. Mice were placed on a warm plate for 2 h after surgery, received a subcutaneous injection of a non-steroidal anti-inflammatory drug (meloxicam, 2 mg/kg) during 3 days, and allowed to recover for 3 weeks before starting behavioral experiments.

### Behavioral experiments

#### Rotarod

4-month mice were trained at accelerating speed (4 - 40 rpm for 5 min), with four sessions per day for three consecutive days and the latency to fall was recorded.

#### Locomotor activity

Mice were placed either in a open-field chamber (50 cm × 50 cm, L x W) for cocaine response or in a 20-cm diameter cylinder for SKF-81297 [SKF] or trihexyphenidyl [THX] response. After 30 minutes, mice were i.p. injected with cocaine (20 mg/kg), SKF (3 mg/kg), or THX (15 mg/kg) and placed back in the chamber for 1 hour. Locomotion was recorded using an overhead digital camera. The distance traveled was measured in 5-min bins using EthoVision software (Noldus, Wageningen, The Netherlands).

#### Conditioned place preference

(CPP) was performed in two compartments of a Y-shaped maze (Imetronic, Pessac, France) with different wall textures and visual cues as follows. (i) Pretest: day-1, mice were placed in the center of the apparatus and allowed to explore freely both compartments for 20 min. The time spent in each compartment was recorded and the preferred and un-preferred compartments deduced for every mouse. (ii) Conditioning: day-2, mice were injected with saline and placed immediately in the un-preferred compartment for 15 min. The next day, they were placed in the other closed compartment after cocaine injection (15 mg/kg). This was repeated twice (3 saline-, 3 cocaine-pairings in total). (iii) Test: time spent in each compartment was measured on day-8 during 20 min. The CPP-score was calculated as the time spent in the cocaine-paired compartment during the test minus the time spent in this compartment during the pre-test.

### Tissue preparation and immunofluorescence

Mice were euthanized, brains rapidly dissected and divided at the midline; one hemisphere was drop-fixed in 40 g/L paraformaldehyde (PFA) for 24 hours. The other hemisphere was flash-frozen using CO_2_ pellets and stored at −80° C. Following PFA fixation, 30-µm-thick sections were cut with a Vibratome (Leica, Wetzlar, Germany). Sections were incubated overnight at 4 °C with primary antibodies, rinsed several times in TBS, and incubated for 45 min with secondary antibody (all antibodies used are in **Table S1**). Nuclei were labelled with DAPI-containing Vectashield (Vector Laboratories, Burlingame, CA, USA). Pyk2 and GFP overall distribution was imaged with a DM6000–2 microscope (Leica). Calbindin and Pyk2 images were acquired with a Leica Confocal SP5-II (63× numerical aperture lens, 5× digital zoom, 1-Airy unit pinhole, 4-frame averaging per *z*-step, *z*-stacks every 2 μm, 1024 × 1024 pixel resolution). Images were analyzed with Icy open source software (https://icy.bioimageanalysis.org) [30].

### Immunobloting

For the analysis of striatal proteins untreated mice were killed by cervical dislocation, striata dissected out, frozen using CO_2_ pellets and stored at −80 °C until use. For pharmacological responses mice were i.p. injected with cocaine (20 mg/kg) or saline and placed in a 43 cm x 27 cm cage. After 10 minutes, mice were euthanized and heads were dipped in liquid nitrogen for 12 seconds. The frozen heads were cut into 210-μm-thick slices with a cryostat, and 10 frozen microdisks (1.4 mm diameter) were punched out bilaterally from the striatum and stored at −80°C until use. Tissue samples were sonicated in 10 g/L SDS and 1 mM sodium orthovanadate in water, and placed at 100 °C for 5 min. Extracts (15 μg protein) were separated by SDS–PAGE and transferred to nitrocellulose membranes (GE Healthcare, Chicago, IL, USA). Membranes were blocked in TBS-T (150 mM NaCl, 20 mM Tris-HCl, pH 7.5, 0.5 ml l^−1^ Tween 20) with 30 g/L BSA. Membranes were incubated overnight at 4 °C with primary antibodies (**Table S1**), washed several times in TBS-T, and incubated with secondary antibodies, which were detected by Odyssey infrared imaging (Li-Cor Inc., Lincoln, NE, USA). For loading control a mouse monoclonal β-actin antibody was used.

### Immunoprecipitation

Mice were i.p. injected with cocaine (20 mg/kg) and placed in a 43 cm x 27 cm cage. After 10 minutes, mice were euthanized and heads dipped in liquid nitrogen for 5 seconds. The striatum was dissected out, lysed by sonication in 250 μl NP-40 lysis buffer [150 mM NaCl, 50 mM Tris-HCl, pH 8.0, 10 mM NaF, 1% NP40 (v/v) supplemented with 1 mM sodium orthovanadate, phosphatase inhibitor (PhosSTOP, Roche, Basel, Switzerland) and protease inhibitor (Complete, Roche)]. Lysates were centrifuged for 20 min at 20,937 g (4°C). Protein A-Sepharose beads (GE Healthcare) were pre-cleared by mixing with 14% Sephacryl S-100 (v/v) (GE Healthcare) and saturated with BSA (25 g/L). The beads were then mixed for 1 hour at 4 °C with rabbit polyclonal anti-Pyk2 antibody (1.7% v/v) (#P3902, Sigma-Aldrich, St. Louis, MO, USA) prior incubation overnight at 4°C with supernatant. The beads were washed three times, resuspended in Laemmli loading buffer, heated at 100 °C for 10 min, and subjected to SDS-PAGE.

### Statistical analysis

Analyses were done using Prism version 6.00 for Windows (GraphPad Software, La Jolla, CA, USA). Data are expressed as means ± SEM. Normal distribution was tested with d’Agostino and Pearson omnibus, Shapiro-Wilk, and Kolmogorov-Smirnov tests. If no difference from normality was detected, statistical analysis was performed using two-tailed Student’s t-test or ANOVA and Holm-Sidak’s post-hoc test. Otherwise non-parametric Mann and Whitney or Kruskal-Wallis’ and Dunn’s tests were used. p < 0.05 was considered as significant.

## Results

### Pyk2 is enriched in ventral D1 SPNs

We used immunohistochemistry to evaluate the regional distribution of Pyk2 in the striatum. Pyk2 immunoreactivity was enriched in the NAc as opposed to the DS (**Figure 1A**), in agreement with a previous study of mRNA distribution [31]. Since Pyk2 striatal immunoreactivity displayed an irregular pattern, we assessed whether its distribution followed the neurochemically-defined patch/matrix compartmentalization [32–34] defined by calbindin, a matrix-enriched protein (**Figures S1A** and **B**). Pyk2 was slightly more expressed in calbindin-positive neurons indicating an enrichment of Pyk2 in matrix compartments (**Figure S1C**). To evaluate the relative expression of Pyk2 in neurons of the direct and indirect pathways, we compared its decrease following conditional deletion. Pyk2 striatal protein levels were decreased by 67 % in Pyk2^f/f;D1::Cre^ mice and by 36% in Pyk2^f/f;A2A::Cre^ mice as compared to matched Pyk2^f/f^ mice (**Figure 1B**), suggesting a higher expression of Pyk2 in D1R-expressing SPNs. Although the wider expression of D1R during the development of the striatum than in the adult [35,36] could lead to a broader developmental action of D1::Cre and subsequent overestimation of the apparent enrichment of Pyk2 in D1 neurons, the conclusion was strengthened by the parallel lower Pyk2 protein decrease in Pyk2^f/f;A2A::Cre^ mice.

**Figure 1:**
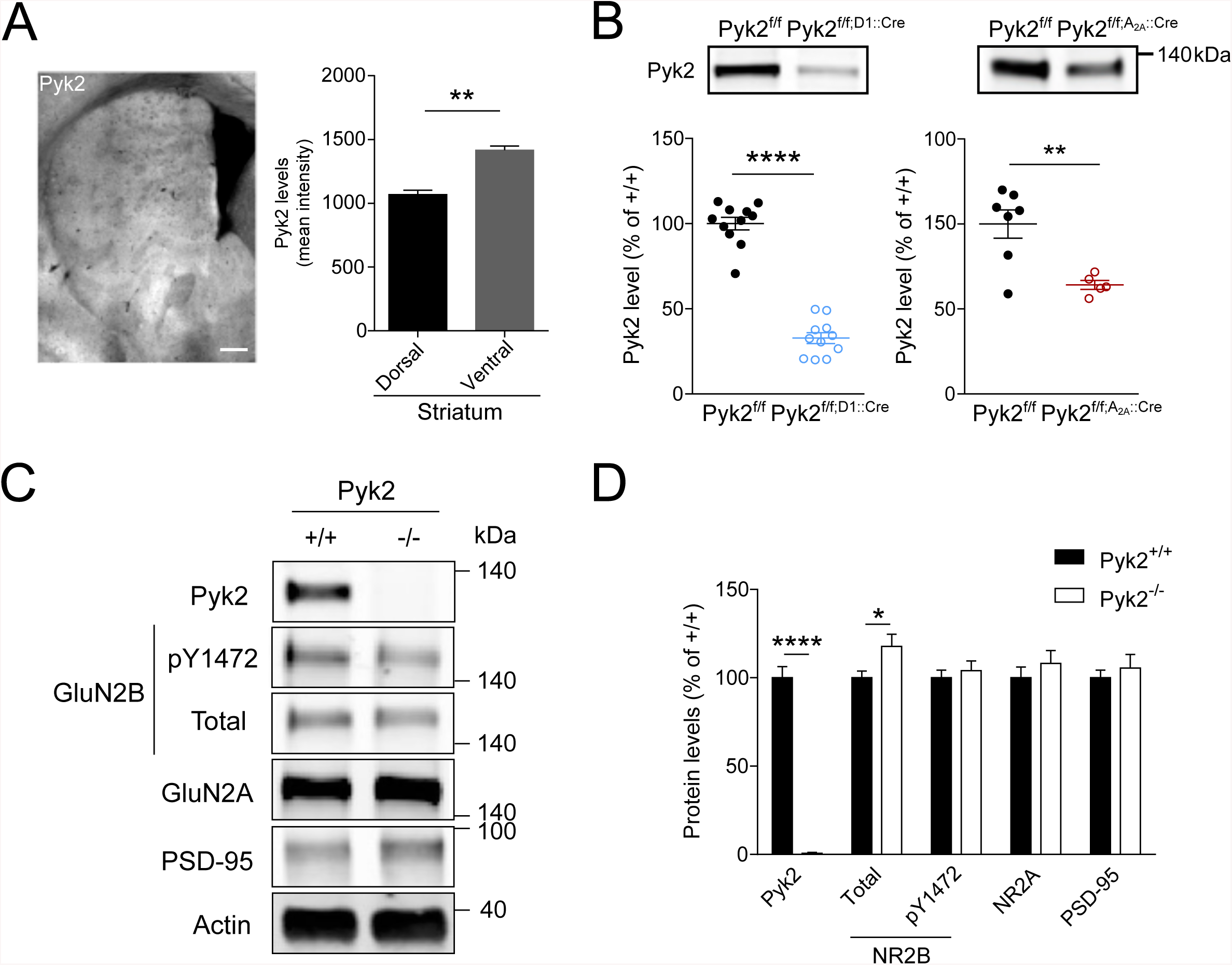
Characterization of Pyk2 expression in the striatum. **a** Left panel, dorsoventral distribution of Pyk2 immunoreactivity in the striatum of wild type mice. Right panel, Pyk2 labelling intensity in the dorsal and ventral striatum was measured in 8 sections from −0.2 to 1.2 mm to bregma, averaged and plotted (3 mice). Scale bar: 300 µm. Unpaired t test, ** p<0.01. **b** Immunoblotting analysis of Pyk2 protein in the striatum of Pyk2^*f/f;D1::Cre*^ and Pyk2^*f/f;A2A::Cre*^, and matched Pyk2^f/f^ control mice. Unpaired t-test, \*\*\*\**p* < 0.0001, \*\**p* < 0.01. **c** Immunoblotting analysis of Pyk2, NMDA receptors subunits, PSD-95 and actin as loading control, in 4-month Pyk2^+/+^ and Pyk2^−/−^ mice. **d** Densitometry quantification of results as in **c**. Data were normalized to actin for each sample and expressed as percentage of wild type mean density. Unpaired t-test, \**p* < 0.05 and \*\*\*\**p* < 0.0001. For number of replicates and detailed statistical analysis, see **Supplementary Table 2**.

### Pyk2 deletion does not alter striatal proteins

In the hippocampus of Pyk2^-/-^ mice, we previously observed decreased levels of several synaptic proteins [15]. In the striatum, neither GluN2A nor PSD-95 levels were altered in Pyk2^-/-^ mice (**Figures 1C** and **D**). GluN2B levels were slightly increased in the striatum of Pyk2^-/-^ mice without change in its phosphorylation. To examine the overall status of SPNs, we measured SPN-enriched proteins, Gαolf [37] and dopamine- and cAMP-regulated neuronal phosphoprotein (DARPP-32) [38]. Their levels were not altered by the mutation (**Figures S2A** and **B**). We finally assessed whether Pyk2 deletion affected a general presynaptic marker synapsin 1 or a dopamine terminals marker tyrosine hydroxylase (TH) in the striatum. We did not observe any change in these protein levels (**Figures S2C** and **D**). These results show that, in contrast to other brain regions, the absence of Pyk2 does not alter basal synaptic or neuronal markers, suggesting regional differences in its function.

### Pyk2 knockout does not impair motor coordination

To study the role of Pyk2 in the striatum, we first assessed motor coordination. Pyk2^+/+^ and Pyk2^-/-^ mice were trained on an accelerating rotarod for 3 days and their motor coordination was evaluated by measuring the time they were able to stay on the rod during each trial. Mice of both genotypes displayed similar performance, improving their ability to stay on the rod trial after trial (**Figure S3A**). Similarly, specific deletion of Pyk2 in the NAc or the DS induced by AAV-mediated Cre expression, or in Pyk2^f/f;D1::Cre^ or Pyk2^f/f;A2A::Cre^ mice had no effect on the rotarod performance (**Figures S3B-E**), showing that motor coordination in this test is not altered in the absence of Pyk2.

### Pyk2 is activated by cocaine and involved in its acute locomotor effects but not in sensitization

Considering the relative enrichment of Pyk2 in the NAc, a region involved in reward response and addiction, we assessed a possible role of Pyk2 in the effects of cocaine. We immunoprecipitated Pyk2 from striatal extracts of saline- or cocaine-injected mice and measured its tyrosine phosphorylation by immunoblotting (**Figure 2A**). Tyrosine phosphorylation of Pyk2 was increased 10 min after cocaine injection, indicating its activation in response to cocaine injection (**Figure 2B**).

**Figure 2:**
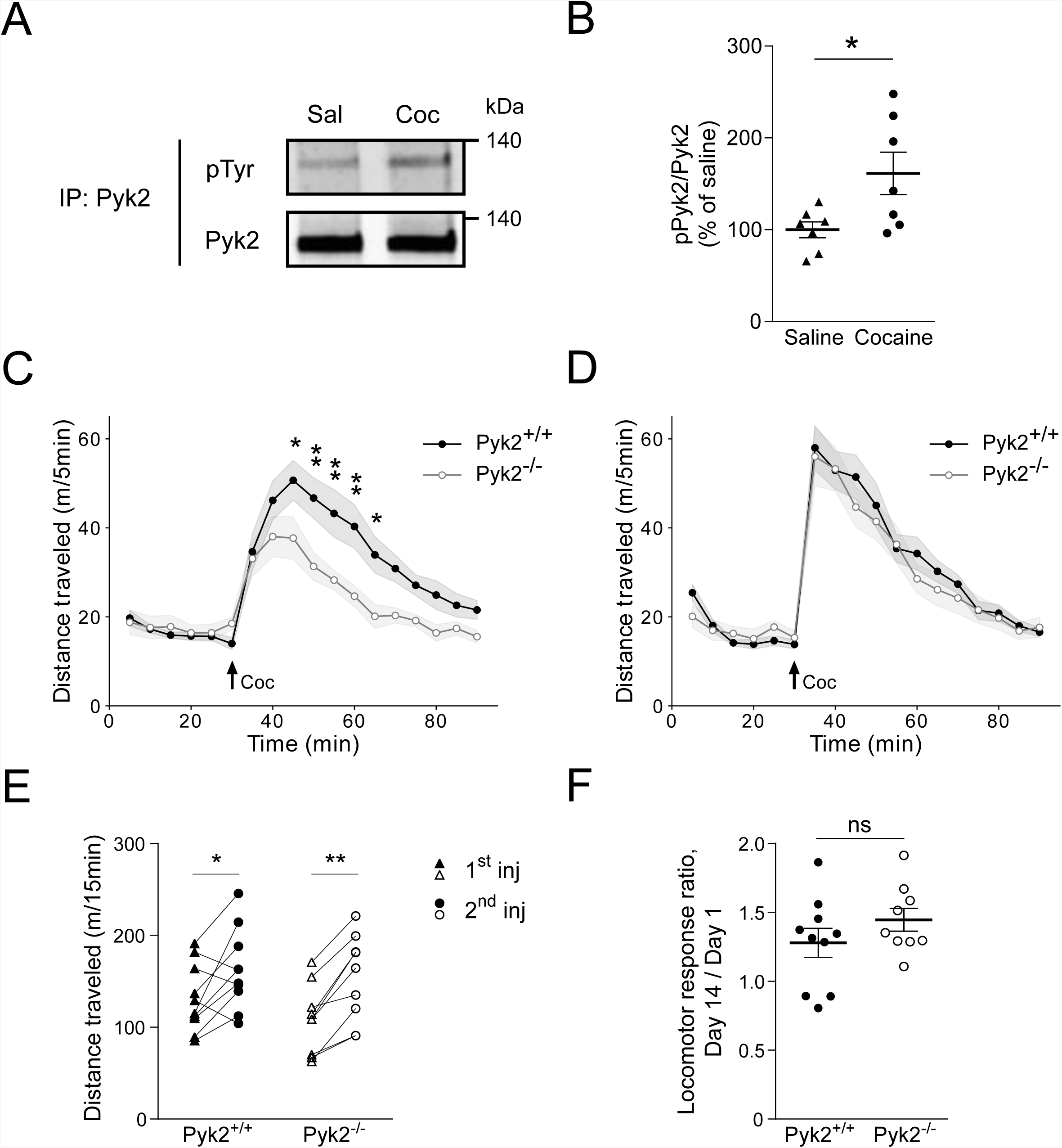
Pyk2 is implicated in acute cocaine responses. **a**, Phospho-Tyr immunoblotting of Pyk2 immunoprecipitated (IP) from striatal extracts of wild-type mice killed 10 min after i.p. injection of saline (Sal) or cocaine (20 mg/kg, Coc). **b**, Densitometry quantification of results as in A. Data are ratios of phospho/total Pyk2 for each sample and expressed as percentage of the mean ratios in saline-treated samples. Unpaired t-test, *p<0.05. **c-d**, Locomotor activity of Pyk2^+/+^ and Pyk2^-/-^ mice after the first (**c**) and the second injection (**d**, 13 days later) of cocaine. Cocaine (20 mg/kg, Coc, arrow) was injected 30 min after mice were placed in the open field. Two-way ANOVA, Sidak’s multiple comparisons post hoc tests, **p < 0.01, *p < 0.05. Values are means ± SEM indicated by a shaded area. **e**-**f**, Locomotor sensitization. **e** Comparison of the distance traveled 0-15 minutes following the first and the second injection of Pyk2^+/+^ and Pyk2^−/−^ mice (from data in **c** and **d**). Two-way ANOVA, Sidak’s multiple comparisons post hoc tests, **p < 0.01, *p < 0.05. **f**, Ratio of the distances traveled 0-15 min after the second (day 14) / first (day 1) cocaine injections, (from data in **e**), in Pyk2^+/+^ and Pyk2^−/−^ mice. Means ± SEM are indicated. Unpaired t-test, ns, not significant. For number of replicates and detailed statistical analysis, see **Supplementary Table 2**.

We then explored a possible role of Pyk2 in the acute response to cocaine by comparing locomotor activity of Pyk2^+/+^ and Pyk2^-/-^ mice after cocaine injection. Basal locomotor activity was similar in the two groups of mice, whereas cocaine-induced hyperlocomotion was decreased in Pyk2^-/-^ mice (**Figure 2C**). Repeated cocaine exposure increases behavioral responses due to a sensitization mechanism [39], which is clearly visible following a single cocaine administration [40]. We therefore injected cocaine a second time, thirteen days later, to assess locomotor sensitization (**Figure 2D**). Locomotor response to the second injection of cocaine was increased in both Pyk2^+/+^ and Pyk2^-/-^ mice (**Figure 2E**) and the 2^nd^ injection/ 1^st^ injection response ratio was not different between the two groups (**Figure 2F**). These results show a role of Pyk2 in the acute cocaine-induced locomotor response but not in its sensitization.

### Pyk2 deficit in the NAc, but not DS, recapitulates the effects of full KO on acute cocaine effects

To identify the striatal region implicated in the effects of Pyk2 deletion, we studied cocaine responses in Pyk2^f/f^ mice bilaterally injected with AAV-GFP-Cre in the NAc or in the DS (**Figure 3**). The position of the tip of the injection needle and the spread of GFP expression was systematically checked on one side of the brain (**Figures 3A** and **B**), the other side being used for immunoblotting. Animals in which the injection was not correctly located were not included in the statistical analysis. Pyk2 immunoreactivity was clearly decreased in the GFP-expressing area (**Figures 3A** and **B**). In mice injected with AAV-GFP-Cre in the NAc, basal locomotor activity was unchanged, but acute locomotor response to cocaine was decreased as compared to mice injected with AAV-GFP (**Figure 3C**). Following a second injection of cocaine at day 14, the locomotor response increased in the two groups of mice and the difference between them was blunted (**Figure 3D**). Sensitization was observed in AAV-GFP and AAV-GFP-Cre mice and the response ratio was not significantly different between the two groups (**Figures S4A** and **3E**). Bilateral injection of AAV-GFP or AAV-GFP-Cre in the DS did not alter the locomotor effects of the first (**Figure 3F**) or second injection of cocaine (**Figures 3G, S4B**, and **3H**). These results provide evidence that the NAc is the region implicated in Pyk2 regulation of cocaine locomotor response.

**Figure 3:**
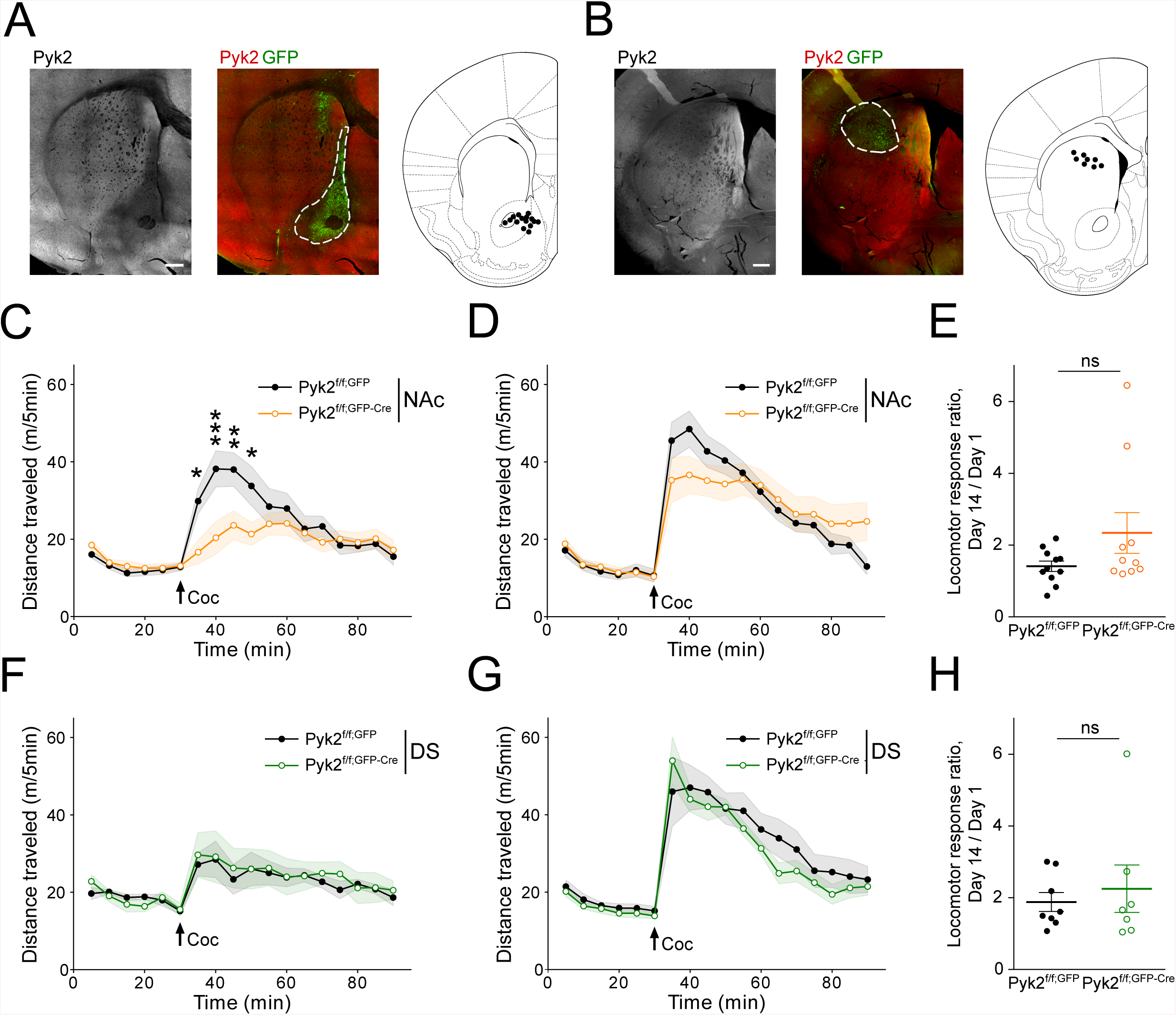
Specific deletion of Pyk2 in the NAc but not in the DS alters the acute locomotor response to cocaine. **a-b** Pyk2^*f/f*^ mice were bilaterally injected in the NAc (**a**) or the DS (**b**) with AAV expressing GFP (Pyk2^*f/f;NAc,GFP*^) or Cre and GFP (Pyk2^*f/f;NAc,GFP-Cre*^). **a**, Anatomical verification of the stereotaxic injections of AAV-GFP-Cre in the NAc: representative Pyk2 immunoreactivity (left panel) and GFP (green) and Pyk2 (red) immunoreactivity colabeling (middle panel). The location of the tips of injecting cannula in all mice used for further analysis is summarized (right panel). **b**, Same as in **a** but for mice receiving injections in the DS. **c-d**, locomotor activity of the mice injected in the NAc (**a**) after the first (**c**) and the second (**d**, 13 days later) injection of cocaine (20 mg/kg, Coc, arrow), as in **Fig. 2c-d**. Two-way ANOVA, AAV effect p<0.001 (**c**), not significant (**d**), Sidak’s multiple comparisons post hoc tests, ***p < 0.001, **p < 0.01, *p < 0.05. **e**, Ratio of the distances traveled 0-15 min after the second (day 14, **d**) / first (day 1, **c**) cocaine injections. Unpaired t-test, not significant, ns. **f-g**, Locomotor activity of the mice injected in the DS (**b**). **f**, Locomotor activity of Pyk2^*f/f;DS,GFP*^ and Pyk2^*f/f;DS,GFP-Cre*^ mice after the first (**f**) and the second (**g**, 13 days later) injection of 20 mg/kg cocaine. Two-way ANOVA, no significant AAV effect. **h**, Ratio of the distances traveled 0-15 min after the second (day 14, **g**) / first (day 1, **f**) cocaine injections in Pyk2^*f/f;DS,GFP*^ and Pyk2^*f/f;DS,GPF-Cre*^ mice. Unpaired t-test, not significant, ns. All values are means ± SEM, indicated by a shaded area in **c, d, f**, and **g**. For number of animals and detailed statistical analysis, see **Supplementary Table 2**.

### Pyk2 deficit in D1, but not A2A SPNs, recapitulates the effects of the full KO on acute cocaine effects

To determine which SPN population was involved in the consequences of Pyk2 deletion, we studied cocaine responses in Pyk2^f/f;D1::Cre^ and Pyk2^f/f;A2A::Cre^ mice (see **Figures 1B** and **4**). Acute locomotor effects of cocaine first injection were decreased in Pyk2^f/f;D1::Cre^ as compared to Pyk2^f/f^ control mice (**Figure 4A**). The response of Pyk2^f/f;D1::Cre^ mice to the second injection of cocaine was still decreased as compared to controls (**Figure 4B**) but a similar sensitization was observed in both genotypes (**Figures S5A** and **4C**). Conversely Pyk2^f/f;A2A::Cre^ mice displayed similar responses as Pyk2^f/f^ mice in response to the first and second injections and comparable sensitization (**Figures 4D-E** and **S5B**). These results indicate that the absence of Pyk2 in D1R-expressing neurons is responsible for the alteration of the locomotor response to cocaine.

**Figure 4:**
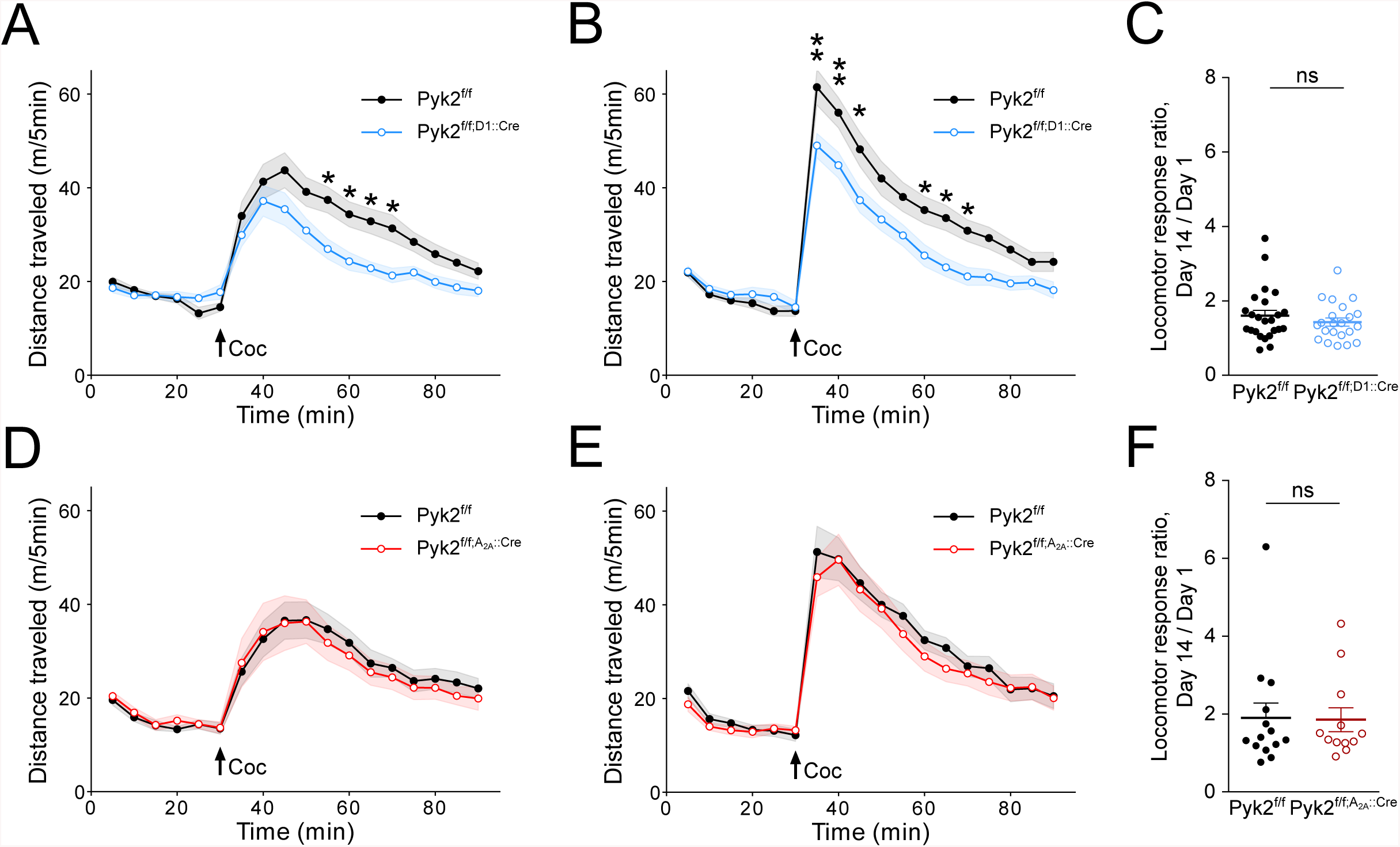
Specific deletion of Pyk2 in D1 but not in A2A neurons alters the locomotor response to cocaine. **a-b**, Locomotor activity of Pyk2^*f/f*^ and Pyk2^*f/f;D1::Cre*^ mice after a first (**a**) and, 13 days later, a second (**b**) injection of cocaine (20 mg/kg, Coc, arrow) as described in the legend to **Fig. 2c-d**. Two-way ANOVA, genotype effect p<0.0001 (**a**), p<0.0001 (**b**), Sidak’s multiple comparisons post hoc tests, **p < 0.01, *p < 0.05. **c**, Ratio of the distances traveled 0-5 min after the second (day 14, **b**) / first (day 1, **a**) cocaine injections in Pyk2^*f/f*^ and Pyk2^*f/f;D1::Cre*^ mice. Unpaired t-test, not significant, ns. **d-e** Same as in **a-b** but in Pyk2^*f/f*^ and Pyk2^*f/f;A2A::Cre*^ mice. Two-way ANOVA, no genotype effect. **f** Ratio of the distances traveled 0-15 min after the second (day 14, **f**) / first (day 1, **d**) cocaine injections in Pyk2^*f/f*^ and Pyk2^*f/f;A2A::Cre*^ mice. Unpaired t-test, ns, not significant. All values are means ± SEM, indicated by a shaded area in **a, b, d**, and **e**. For number of animals and detailed statistical analysis, see **Supplementary Table 2**.

### Pyk2 knockout has minor effects on cocaine-induced signaling and does not alter conditioned place preference

Cocaine injection was reported to increase GluN2B phosphorylation at Tyr-1472 and to activate SFK, thus contributing to ERK activation [8]. To evaluate the role of Pyk2 in this response we compared the effects of cocaine on Tyr1472-GluN2B phosphorylation in Pyk2^f/f;D1::Cre^ and Pyk2^f/f^ mice. In Pyk2-mutant mice, 10 min after cocaine injection, the increase in pTyr1472-GluN2B was blunted and not significant (**Figures S6A** and **B**). This minor alteration may reflect the direct or indirect role of Pyk2 in the phosphorylation of the NMDA receptor. However, the absence of Pyk2 in D1 SPNs did not prevent the activation of ERK (**Figures S6A** and **C**), a step critical for long-term effects of cocaine [7].

We then evaluated the rewarding properties of cocaine in Pyk2 mutant mice, which involve D1 receptors [41–43]. We measured cocaine-CPP in wild-type and mutant mice (**Figure S7**). The time spent in the cocaine-paired arm after conditioning was similarly increased in Pyk2^-/-^ and Pyk2^+/+^ mice (**Figure S7A**). The CPP score did not differ between Pyk2^-/-^ and Pyk2^+/+^ mice (**Figure S7B**), demonstrating that mice underwent efficient CPP in the absence of Pyk2. We also examined whether specific deletion of Pyk2 in the NAc or DS, or in D1R- or A2AR-expressing SPNs had any effect on CPP (**Figure S7C-F**). CPP scores were not modified in any of the mutant groups as compared to their respective controls, confirming the lack of involvement of Pyk2 in CPP. This result is in agreement with the observation that in the absence of Pyk2, ERK signaling pathway, which plays a critical role in cocaine-CPP [44], was still activated.

### Pyk2 is involved in the acute locomotor effects of a D1R agonist but not a cholinergic antagonist

Since the consequences of the absence of Pyk2 on acute locomotor effects of cocaine appeared to result from Pyk2 deficit in D1 neurons, we examined whether effects of D1R stimulation were altered. We used SKF-81297, a D1R selective agonist, known to increase locomotor activity [45–47]. The acute locomotor response to SKF-81297 was slightly reduced in Pyk2^f/f;D1::Cre^ mice as compared to Pyk2^f/f^ mice (**Figure 5A**). As reported in previous studies [48–50], a sensitization of the response was observed following the second injection in wild-type and mutant mice (**Figure 5B**). Interestingly, the first and second injection of SKF-81297 both had a biphasic effect on locomotor activity, not previously reported to our knowledge. We analyzed separately the sensitization during the first 10 minutes following injection (**Figures 5C** and **D**) and from 10 to 40 minutes (**Figures 5E** and **F**). Sensitization was significant for these two different phases without difference between Pyk2^f/f;D1::Cre^ and Pyk2^f/f^ mice (**Figures 5-F**). To determine whether Pyk2 could decrease any kind of drug-induced locomotor activity, we tested the effects of trihexyphenidine (THX), an anticholinergic substance that increases locomotor activity [51]. We found no difference in locomotor activity between Pyk2^f/f;D1::Cre^ and Pyk2^f/f^ mice following the first or second injection of THX (**Figure S8A** and **B**). The responses to the two injections of THX were similar without locomotor sensitization in either genotype (**Figure S8C**). These results indicate that Pyk2 is selectively involved in the locomotor activity induced by stimulation of D1 receptors.

**Figure 5:**
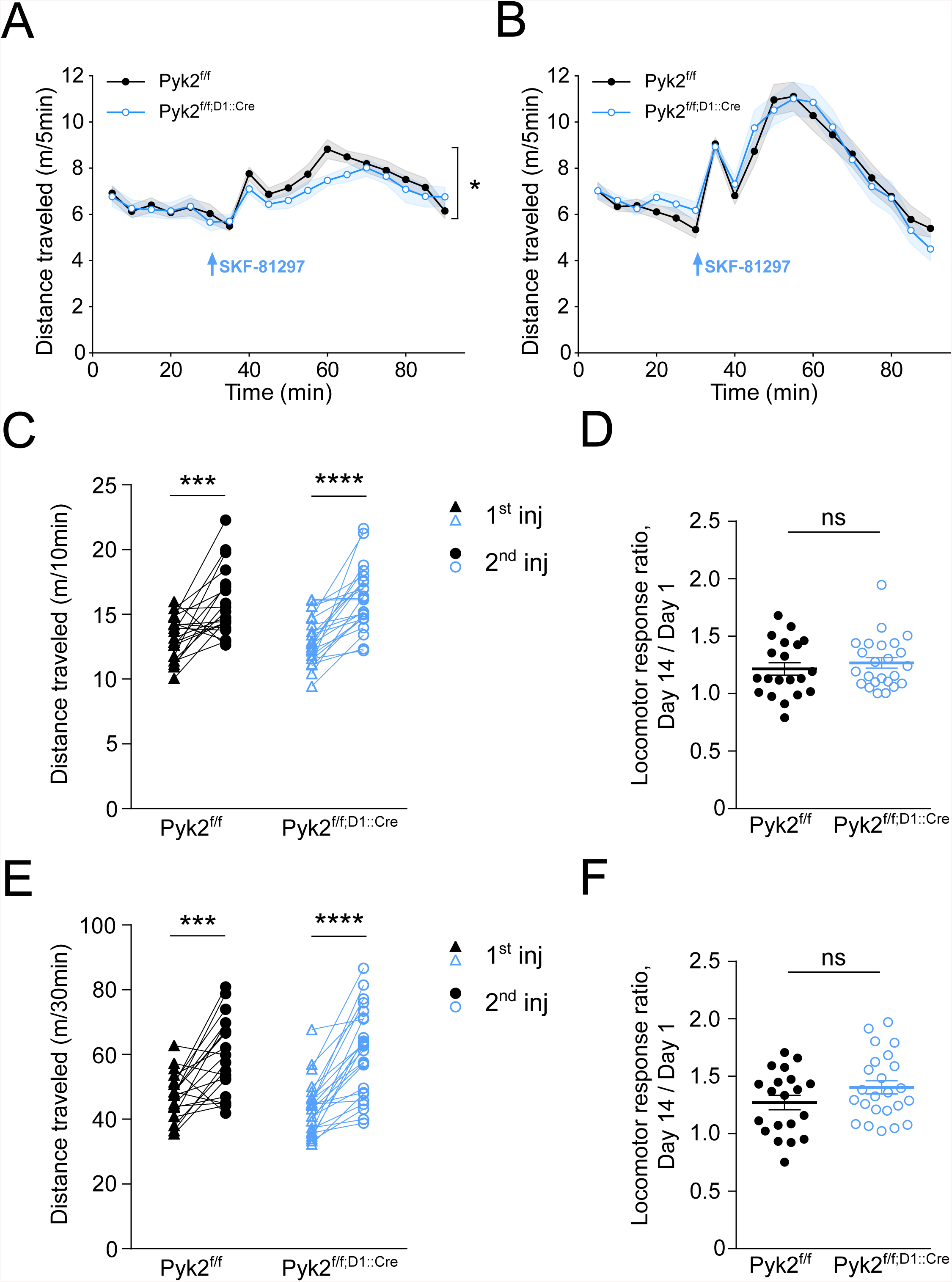
Specific deletion of Pyk2 in D1 neurons alters the acute locomotor response to a D1R agonist, SKF-81297. **a-b**, Locomotor activity of Pyk2^*f/f*^ and Pyk2^*f/f;D1::Cre*^ mice after a first (**a**) and a second (**b**, 13 days later) injection of SKF-81297 (SKF). SKF (3 mg/kg, arrow) was injected 30 min after mice were placed in the open field. Two-way ANOVA, genotype effect, p = 0.016 (**a**), not significant (**b**). **c**-**d**, Sensitization of the response during the first 10 min after SKF injection (data from **a** and **b**). **c**, Distance traveled during the first 10 minutes following the first and the second SKF injections in Pyk2^*f/f*^ and Pyk2^*f/f;D1::Cre*^ mice. Two-way ANOVA, Sidak’s multiple comparisons post hoc test, ****p<0.0001, ***p < 0.001. **d**, Ratios of the distances traveled 0-15 min after the second (day 14, **b**) / first (day 1, **a**) and SKF injections in Pyk2^*f/f*^ and Pyk2^*f/f;D1::Cre*^ mice. Unpaired t-test, ns, not significant. **e**-**f**, Same analyses as in **c** and **d**, but for the distance traveled 15 to 40 minutes after SKF injection (data from **a** and **b**). **a-f**, Values are means ± SEM, indicated by a shaded area in **a** and **b**. For number of animals and detailed statistical analysis, see **Supplementary Table 2**.

## Discussion

In this study, we show the involvement of the non-receptor tyrosine kinase Pyk2 in the acute locomotor response to cocaine. We demonstrate that complete knockout of Pyk2 or its specific deletion in the NAc or in D1R-expressing neurons decreases acute cocaine-induced hyperlocomotion. Acute locomotor response to a selective D1 agonist but not to a cholinergic antagonist was also decreased. These results indicate a role of Pyk2 located in D1R-expressing neurons of the NAc in the D1-induced acute locomotor response. In contrast no alteration was observed in locomotor sensitization or cocaine-CPP.

Our study of the expression pattern of Pyk2 in the striatum confirmed its enrichment of Pyk2 in the ventral part of the striatum, previously reported [31]. We also provide evidence for a relative enrichment in D1R-expressing SPNs which form the ‘direct pathway’ [52] and in the matrix compartment of the striatum which plays a specific yet incompletely characterized role in striatal function [53]. The dorsal striatum is implicated in the control of movement and the learning process of skilled task [54,55]. We assessed motor coordination of mice with accelerating rotarod training [56], which can be impaired by alteration of various brain areas, including basal ganglia, motor cortex, and cerebellum [57]. We did not observe any difference between control and Pyk2 mutant mice, suggesting that Pyk2 is not necessary for the function of the dorsal striatum where it is relatively less expressed. Pyk2 is more expressed in the NAc, which is a key component in brain circuitry underlying drug-evoked behaviors [58,59]. We demonstrated that tyrosine phosphorylation of Pyk2 in the striatum was increased following a single injection of cocaine. Pyk2 is activated by Ca^2+^-induced autophosphorylation of Tyr-402, followed by recruitment of SFKs which in turn phosphorylate Pyk2 on other tyrosine residues, although in some cells activation of SFKs can be the triggering mechanism (see ref. [17] for a review). Cocaine injection in naïve mice increases the concentration of calcium in D1 SPNs [60] providing a possible basis for Pyk2 activation. In addition, cocaine-induced activation of Fyn has been reported in the striatum and proposed to be involved in NMDA receptors regulation and their synergism with D1R to activate the ERK pathway [8]. Although this could suggest a role of Pyk2 in ERK activation, ERK phosphorylation was still observed in Pyk2-deficient mice, underlining the redundancy in the mechanisms controlling ERK activation.

Concerning the functional consequences of Pyk2 knock-out, we found that the acute cocaine locomotor response was decreased, whereas locomotor sensitization and CPP, which are known to involve synaptic plasticity and ERK activation [7], were not altered. The impaired acute locomotor response was also observed in mice specifically lacking Pyk2 in the NAc or in D1R-expressing neurons, but not in those devoid of Pyk2 in the DS or in A_2A_R-expressing neurons. This combination of results implicates an alteration induced by the absence of Pyk2 in the D1R-expressing neurons of the NAc. This conclusion is supported by the role of these neurons in the acute locomotor effects of psychostimulants [61–63]. A decreased locomotor response without alteration in locomotor sensitization or CPP was also observed in G*α*olf heterozygous (*Gnal*^+/−^) mutant mice in which cAMP production is decreased [64]. These similarities could indicate a role for Pyk2 in D1R/G*α*olf/cAMP pathway modulation, a hypothesis supported by slightly lower locomotor response to the D1 agonist SKF-81297 in Pyk2^f/f;D1::Cre^ mice. However we did not find any change in G*α*olf levels in the mutant mice. The only effect we observed was a blunting of GluN2B receptor phosphorylation at Tyr1472, in good agreement with the role of Pyk2 in the phosphorylation of the NMDAR [11], but unlikely to explain by itself the effects on locomotor activity. Therefore we hypothesize that Pyk2 is also implicated in the regulation of other signaling mechanisms important for D1-mediated regulation of locomotor activity. Further work will be necessary to explore this hypothesis and in particular whether Pyk2 modulates the production of cAMP in response to D1R stimulation. Our study clearly identifies the significant but circumscribed contribution of Pyk2 in NAC D1 neurons to the acute locomotor effects of cocaine. It raises the question of its role in other dopamine-mediated actions and opens novel avenues for investigating signaling mechanisms in striatal neurons.

## Funding and Disclosure

BdP was supported in part by the *Fondation pour la Recherche Médicale* (FRM). AG is now a Ramón y Cajal fellow (RYC-2016-19466). JAG lab was supported in part by Inserm, *Sorbonne Université*, the FRM, ERC AdG-2009, a KAUST collaborative grant to JAG and S Arold, and the Bio-Psy (Biology for Psychiatry) laboratory of excellence. Relevant experiments were carried out at the IFM *Rodent breeding and phenotyping* and *Cell and Tissue Imaging* facilities. The *Cell and tissue imaging* facility at the IFM benefited from supports of *Espoir en Tête/Fondation pour la Recherche sur le Cerveau, Région Ile-de-France*, and Inserm.

The authors declare no competing financial interests

## Acknowledgements

The authors thank the late Paul Greengard (The Rockefeller University) for the gift of antibodies for synapsin 1, DARPP-32 and phosphoT75-DARPP-32, and Alban de Kerchove d’Exaerde (*Université libre de Bruxelles*) for providing Adora2a::Cre mice, Manuel Mameli (University of Lausanne) and Kristina Valentinova (University of Bern) for their help when they were at the IFM and for critical reading of the manuscript. We thank Denis Hervé for expert advice.

