## Supplementary material for "Pyk2 in D1 receptor-expressing neurons of the nucleus accumbens modulates the acute locomotor effects of cocaine": Table S1

| Antibody table | | | | |
|---|---|---|---|---|
| **Antibody** | **Species** | **Dilution for WB** | **Dilution for IF** | **Company (reference number)** |
| Pyk2 | Rabbit | 1:1000 | 1:500 | Sigma-Aldrich (#P3902) |
| PSD-95 | Rabbit | 1:1000 | - | Cell Signaling Technology (#3450) |
| phosphoY1472-GluN2B | Rabbit | 1:1000 | - | Cayman chemical (#10009761) |
| GluN2B | Rabbit | 1:1000 | - | Merck KGaA (#06-600) |
| GluN2A | Rabbit | 1:1000 | - | Merck KGaA (#05-901R) |
| Gαolf | Rabbit | 1:1000 | - | produced as described [30] |
| synapsin 1 | Rabbit | 1:500 | - | gift from Pr. Greengard (G-486) |
| tyrosine hydroxylase | Rabbit | 1:1000 | - | Merck KGaA (#AB1541) |
| STEP | Rabbit | 1:1000 | - | Cell Signaling Technology (#9069) |
| phosphoS845-GluA1 | Rabbit | 1:1000 | - | Abcam (#ab3901) |
| phosphoT202/Y204-Erk1/2 | Rabbit | 1:1000 | - | Cell Signaling Technology (#9101) |
| Erk1/2 | Rabbit | 1:1000 | - | Cell Signaling Technology (#9102) |
| Calbindin D28k | Mouse | - | 1:1000 | Swant (#300) |
| Pyk2 | Mouse | 1:1000 | - | Cell Signaling Technology (#3480) |
| DARPP-32 | Mouse | 1:5000 | - | gift from Pr. Greengard (m6Ab) |
| phosphoT75-DARPP-32 | Mouse | 1:1000 | - | gift from Pr. Greengard (RU911) |
| phosphotyrosine | Mouse | 1:1000 | - | Cell Signaling Technology (#05-947) |
| β-actin | Mouse | 1:5000 | - | Sigma-Aldrich (#A5441) |
| tyrosine hydroxylase | Chicken | 1:1000 | - | Aves Labs (#TH) |
| GFP | Chicken | - | 1:500 | Thermo Fisher (#A10262) |
| anti-rabbit IgG DyLight™ 680 | Goat | 1:10000 | - | Rockland Immunochemicals (#611-144-002) |
| anti-rabbit IgG DyLight™ 800 | Goat | 1:10000 | - | Rockland Immunochemicals (#611-145-002) |
| anti-mouse IgG DyLight™ 680 | Goat | 1:10000 | - | Rockland Immunochemicals (#610-144-002) |
| anti-mouse IgG DyLight™ 800 | Goat | 1:10000 | - | Rockland Immunochemicals (#610-145-002) |
| anti-chicken IgG DyLight™ 800 | Goat | 1:10000 | - | Rockland Immunochemicals (#603-145-002) |
| anti-Rabbit IgG Alexa Fluor 555 | Donkey | - | 1:400 | Thermo Fisher (#A-31572) |
| anti-Mouse IgG Alexa Fluor 488 | Goat | - | 1:400 | Thermo Fisher (#A-11001) |
| anti-Chicken IgG Alexa Fluor 488 | Goat | - | 1:400 | Thermo Fisher (#A-11039) |
