## Supplementary material for "Pyk2 in D1 receptor-expressing neurons of the nucleus accumbens modulates the acute locomotor effects of cocaine": Table S2

| Figure | Group | Mean | SEM | Comparison | Statistical analysis | DF | Test value | *P* value | n |
|---|---|---|---|---|---|---|---|---|---|
| 1A | Dorsal | 1069 | 33.22 | | | | | | 3 mice |
| | Ventral | 1416 | 33.36 | | | | | | 3 mice |
| | | | | Dorsal vs Ventral | Two-tailed unpaired t-test | 4 | t = 7.37 | **0.0018** | |
| 1B left panel | $Pyk2^{f/f}$ | 100 | 3.70 | | | | | | 11 mice |
| | $Pyk2^{f/f;D1::Cre}$ | 32.76 | 3.20 | | | | | | 11 mice |
| | | | | $Pyk2^{f/f}$ vs $Pyk2^{f/f;D1::Cre}$ | Two-tailed unpaired t-test | 20 | t = 13.75 | **< 0.0001** | |
| 1B right panel | $Pyk2^{f/f}$ | 100 | 8.31 | | | | | | 7 mice |
| | $Pyk2^{f/f;A2A::Cre}$ | 64.13 | 2.63 | | | | | | 5 mice |
| | | | | $Pyk2^{f/f}$ vs $Pyk2^{f/f;A2A::Cre}$ | Two-tailed unpaired t-test | 10 | t = 3.515 | **0.0056** | |
| 1D | Pyk2 - $Pyk2^{+/+}$ | 100 | 6.36 | | | | | | 6 mice |
| | Pyk2 - $Pyk2^{-/-}$ | 0.72 | 0.45 | | | | | | 6 mice |
| | | | | Pyk2 - $Pyk2^{+/+}$ vs $Pyk2^{-/-}$ | Two-tailed unpaired t-test | 10 | t = 15.57 | **< 0.0001** | |
| | NR2B - $Pyk2^{+/+}$ | 100 | 3.76 | | | | | | 15 mice |
| | NR2B - $Pyk2^{-/-}$ | 117.79 | 6.92 | | | | | | 16 mice |
| | | | | NR2B - $Pyk2^{+/+}$ vs $Pyk2^{-/-}$ | Two-tailed unpaired t-test | 29 | t = 2.22 | **0.0346** | |
| | pY1472NR2B - $Pyk2^{+/+}$ | 100 | 4.35 | | | | | | 15 mice |
| | pY1472NR2B - $Pyk2^{-/-}$ | 104.04 | 5.5 | | | | | | 16 mice |
| | | | | pY1472NR2B - $Pyk2^{+/+}$ vs $Pyk2^{-/-}$ | Two-tailed unpaired t-test | 29 | t = 0.57 | 0.5721 | |
| | NR2A - $Pyk2^{+/+}$ | 100 | 6.12 | | | | | | 6 mice |
| | NR2A - $Pyk2^{-/-}$ | 108.23 | 7.24 | | | | | | 6 mice |
| | | | | NR2A - $Pyk2^{+/+}$ vs $Pyk2^{-/-}$ | Two-tailed unpaired t-test | 10 | t = 0.87 | 0.4058 | |
| | PSD-95 - $Pyk2^{+/+}$ | 100 | 4.37 | | | | | | 6 mice |
| | PSD-95 - $Pyk2^{-/-}$ | 105.61 | 7.53 | | | | | | 8 mice |
| | | | | PSD-95 - $Pyk2^{+/+}$ vs $Pyk2^{-/-}$ | Two-tailed unpaired Mann Whitney test | | U = 24 | > 0.9999 | |
| 2B | Saline | 100 | 8.74 | | | | | | 7 mice |
| | Cocaine | 161.3 | 23.05 | | | | | | 7 mice |
| | | | | Saline vs Cocaine | Two-tailed unpaired t-test | 12 | t = 2.49 | **0.0286** | |
| 2C | $Pyk2^{+/+}$ | | | | | | | | 10 mice |
| | $Pyk2^{-/-}$ | | | | | | | | 9 mice |
| | | | | | Two-way ANOVA: | | | | |
| | | | | | Interaction | 17 | F = 2.44 | **0.0014** | |
| | | | | | Time | 17 | F = 20.43 | **< 0.0001** | |
| | | | | | Genotype | 1 | F = 39.05 | **< 0.0001** | |
| 2D | $Pyk2^{+/+}$ | | | | | | | | 10 mice |
| | $Pyk2^{-/-}$ | | | | | | | | 9 mice |
| | | | | | Two-way ANOVA: | | | | |
| | | | | | Interaction | 17 | F = 0.43 | 0.9769 | |
| | | | | | Time | 17 | F = 40.18 | **< 0.0001** | |
| | | | | | Genotype | 1 | F = 1.59 | 0.208 | |

| | | | | | | | | | |
|---|---|---|---|---|---|---|---|---|---|
| 2E | 1st injection - $Pyk2^{+/+}$ | 13147 | 1163.53 | | | | | | 10 mice |
| | 2nd injection - $Pyk2^{+/+}$ | 16230 | 1383.59 | | | | | | 9 mice |
| | 1st injection - $Pyk2^{-/-}$ | 10885 | 1264.40 | | | | | | 10 mice |
| | 2nd injection - $Pyk2^{-/-}$ | 15390 | 1567.73 | | | | | | 9 mice |
| | | | | | Two-way ANOVA: | | | | |
| | | | | | Interaction | 1 | F = 0.90 | 0.3572 | |
| | | | | | Genotype | 1 | F = 0.78 | 0.3886 | |
| | | | | | Injection | 1 | F = 25.49 | **< 0.0001** | |
| | | | | | Subjects (matching) | 17 | F = 5.44 | **0.0005** | |
| | | | | $Pyk2^{+/+}$: 1st inj vs 2nd inj | Sidak's multiple comparisons test | 17 | t = 2.98 | **0.0167** | |
| | | | | $Pyk2^{-/-}$: 1st inj vs 2nd inj | Sidak's multiple comparisons test | 17 | t = 4.13 | **0.0014** | |
| 2F | $Pyk2^{+/+}$ | 1.28 | 0.1 | | | | | | 10 mice |
| | $Pyk2^{-/-}$ | 1.45 | 0.08 | | | | | | 9 mice |
| | | | | $Pyk2^{+/+}$ vs $Pyk2^{-/-}$ | Two-tailed unpaired t-test | 17 | t = 1.24 | 0.2315 | |
| 3C | $Pyk2^{f/f;NAc,GFP}$ | | | | | | | | 11 mice |
| | $Pyk2^{f/f;NAc,GFP-Cre}$ | | | | | | | | 10 mice |
| | | | | | Two-way ANOVA: | | | | |
| | | | | | Interaction | 17 | F = 2.85 | **0.0002** | |
| | | | | | Time | 17 | F = 10.15 | **< 0.0001** | |
| | | | | | AAV type | 1 | F = 12.94 | **0.0004** | |
| 3D | $Pyk2^{f/f;NAc,GFP}$ | | | | | | | | 11 mice |
| | $Pyk2^{f/f;NAc,GFP-Cre}$ | | | | | | | | 10 mice |
| | | | | | Two-way ANOVA: | | | | |
| | | | | | Interaction | 17 | F = 1.49 | 0.0942 | |
| | | | | | Time | 17 | F = 22.16 | **< 0.0001** | |
| | | | | | AAV type | 1 | F = 0.02 | 0.9015 | |
| 3E | $Pyk2^{f/f;NAc,GFP}$ | 1.41 | 0.14 | | | | | | 11 mice |
| | $Pyk2^{f/f;NAc,GFP-Cre}$ | 2.34 | 0.57 | | | | | | 10 mice |
| | | | | $Pyk2^{f/f;NAc,GFP}$ vs $Pyk2^{f/f;NAc,GFP-Cre}$ | Two-tailed unpaired Mann-Whitney test | | U = 39 | 0.2785 | |
| 3F | $Pyk2^{f/f;DS,GFP}$ | | | | | | | | 8 mice |
| | $Pyk2^{f/f;DS,GFP-Cre}$ | | | | | | | | 7 mice |
| | | | | | Two-way ANOVA: | | | | |
| | | | | | Interaction | 17 | F = 0.16 | > 0.9999 | |
| | | | | | Time | 17 | F = 3.00 | **< 0.0001** | |
| | | | | | AAV type | 1 | F = 0.55 | 0.4586 | |
| 3G | $Pyk2^{f/f;DS,GFP}$ | | | | | | | | 8 mice |
| | $Pyk2^{f/f;DS,GFP-Cre}$ | | | | | | | | 7 mice |
| | | | | | Two-way ANOVA: | | | | |
| | | | | | Interaction | 17 | F = 0.78 | 0.7197 | |
| | | | | | Time | 17 | F = 22.93 | **< 0.0001** | |
| | | | | | AAV type | 1 | F = 7.93 | **0.0053** | |
| 3H | $Pyk2^{f/f;DS,GFP}$ | 1.88 | 0.26 | | | | | | 8 mice |
| | $Pyk2^{f/f;DS,GFP-Cre}$ | 2.25 | 0.66 | | | | | | 7 mice |
| | | | | $Pyk2^{f/f;DS,GFP}$ vs $Pyk2^{f/f;DS,GFP-Cre}$ | Two-tailed unpaired Mann-Whitney test | | U = 27 | 0.9203 | |

| | | | | | | | | | |
|---|---|---|---|---|---|---|---|---|---|
| 4A | Pyk2$^{f/f}$ | | | | | | | | 24 mice |
| | Pyk2$^{f/f;D1::Cre}$ | | | | | | | | 22 mice |
| | | | | | Two-way ANOVA: | | | | |
| | | | | | Interaction | 17 | F = 1.73 | **0.0329** | |
| | | | | | Time | 17 | F = 24.04 | **< 0.0001** | |
| | | | | | Genotype | 1 | F = 35.32 | **< 0.0001** | |
| 4B | Pyk2$^{f/f}$ | | | | | | | | 24 mice |
| | Pyk2$^{f/f;D1::Cre}$ | | | | | | | | 22 mice |
| | | | | | Two-way ANOVA: | | | | |
| | | | | | Interaction | 17 | F = 2.38 | **0.0014** | |
| | | | | | Time | 17 | F = 52.38 | **< 0.0001** | |
| | | | | | Genotype | 1 | F = 47.46 | **< 0.0001** | |
| 4C | Pyk2$^{f/f}$ | 1.60 | 0.14 | | | | | | 24 mice |
| | Pyk2$^{f/f;D1::Cre}$ | 1.43 | 0.11 | | | | | | 22 mice |
| | | | | Pyk2$^{+/+}$ vs Pyk2$^{f/f;D1::Cre}$ | Two-tailed unpaired t-test | 44 | t = 0.95 | 0.3471 | |
| 4D | Pyk2$^{f/f}$ | | | | | | | | 14 mice |
| | Pyk2$^{f/f;A2A::Cre}$ | | | | | | | | 12 mice |
| | | | | | Two-way ANOVA: | | | | |
| | | | | | Interaction | 17 | F = 0.17 | > 0.9999 | |
| | | | | | Time | 17 | F = 14.73 | **< 0.0001** | |
| | | | | | Genotype | 1 | F = 0.44 | 0.5077 | |
| 4E | Pyk2$^{f/f}$ | | | | | | | | 14 mice |
| | Pyk2$^{f/f;A2A::Cre}$ | | | | | | | | 12 mice |
| | | | | | Two-way ANOVA: | | | | |
| | | | | | Interaction | 17 | F = 0.22 | 0.9997 | |
| | | | | | Time | 17 | F = 38.01 | **< 0.0001** | |
| | | | | | Genotype | 1 | F = 2.85 | 0.0924 | |
| 4F | Pyk2$^{f/f}$ | 1.90 | 0.38 | | | | | | 14 mice |
| | Pyk2$^{f/f;A2A::Cre}$ | 1.85 | 0.31 | | | | | | 12 mice |
| | | | | Pyk2$^{+/+}$ vs Pyk2$^{f/f;A2A::Cre}$ | Two-tailed unpaired Mann-Whitney test | | U = 79 | 0.8101 | |
| 5A | Pyk2$^{f/f}$ | | | | | | | | 20 mice |
| | Pyk2$^{f/f;D1::Cre}$ | | | | | | | | 27 mice |
| | | | | | Two-way ANOVA: | | | | |
| | | | | | Interaction | 17 | F = 0.71 | 0.7985 | |
| | | | | | Time | 17 | F = 9.14 | **< 0.0001** | |
| | | | | | Genotype | 1 | F = 5.80 | **0.0163** | |
| 5B | Pyk2$^{f/f}$ | | | | | | | | 20 mice |
| | Pyk2$^{f/f;D1::Cre}$ | | | | | | | | 24 mice |
| | | | | | Two-way ANOVA: | | | | |
| | | | | | Interaction | 17 | F = 0.47 | 0.9648 | |
| | | | | | Time | 17 | F = 26.32 | **< 0.0001** | |
| | | | | | Genotype | 1 | F = 0.36 | 0.5485 | |

| | | | | | | | | | |
|---|---|---|---|---|---|---|---|---|---|
| 5C | 1st injection - Pyk2$^{f/f}$ | 13.25 | 0.38 | | | | | | 20 mice |
| | 2nd injection - Pyk2$^{f/f}$ | 15.86 | 0.59 | | | | | | 24 mice |
| | 1st injection - Pyk2$^{f/f;D1::Cre}$ | 12.97 | 0.38 | | | | | | 20 mice |
| | 2nd injection - Pyk2$^{f/f;D1::Cre}$ | 16.24 | 0.49 | | | | | | 24 mice |
| | | | | | Two-way ANOVA: | | | | |
| | | | | | Interaction | 1 | F = 0.60 | 0.4416 | |
| | | | | | Genotype | 1 | F = 0.01 | 0.9188 | |
| | | | | | Injection | 1 | F = 48.70 | **< 0.0001** | |
| | | | | | Subjects (matching) | 42 | F = 1.44 | 0.1197 | |
| | | | | Pyk2$^{f/f}$: 1st inj vs 2nd inj | Sidak's multiple comparisons test | 42 | t = 4.20 | **0.0003** | |
| | | | | Pyk2$^{f/f;D1::Cre}$: 1st inj vs 2nd inj | Sidak's multiple comparisons test | 42 | t = 5.75 | **< 0.0001** | |
| 5D | Pyk2$^{f/f}$ | 1.22 | 0.05 | | | | | | 20 mice |
| | Pyk2$^{f/f;D1::Cre}$ | 1.27 | 0.05 | | | | | | 24 mice |
| | | | | Pyk2$^{+/+}$ vs Pyk2$^{f/f;D1::Cre}$ | Two-tailed unpaired t-test | 42 | t = 0.75 | 0.4589 | |
| 5E | 1st injection - Pyk2$^{f/f}$ | 47.24 | 1.58 | | | | | | 20 mice |
| | 2nd injection - Pyk2$^{f/f}$ | 59.13 | 2.64 | | | | | | 24 mice |
| | 1st injection - Pyk2$^{f/f;D1::Cre}$ | 43.38 | 1.72 | | | | | | 20 mice |
| | 2nd injection - Pyk2$^{f/f;D1::Cre}$ | 60.27 | 2.76 | | | | | | 24 mice |
| | | | | | Two-way ANOVA: | | | | |
| | | | | | Interaction | 1 | F = 1.91 | 0.1745 | |
| | | | | | Genotype | 1 | F = 0.26 | 0.6103 | |
| | | | | | Injection | 1 | F = 63.19 | **< 0.0001** | |
| | | | | | Subjects (matching) | 42 | F = 2.14 | **0.0077** | |
| | | | | Pyk2$^{f/f}$: 1st inj vs 2nd inj | Sidak's multiple comparisons test | 42 | t = 2.98 | **0.0167** | |
| | | | | Pyk2$^{f/f;D1::Cre}$: 1st inj vs 2nd inj | Sidak's multiple comparisons test | 42 | t = 4.13 | **0.0014** | |
| 5F | Pyk2$^{f/f}$ | 1.27 | 0.06 | | | | | | 20 mice |
| | Pyk2$^{f/f;D1::Cre}$ | 1.40 | 0.06 | | | | | | 24 mice |
| | | | | Pyk2$^{+/+}$ vs Pyk2$^{f/f;D1::Cre}$ | Two-tailed unpaired t-test | 42 | t = 1.56 | 0.1268 | |
| S1C | Matrix | 2910 | 40.9 | | | | | | 20 regions (from 3 mice) |
| | Patch | 2396 | 58.53 | | | | | | 20 regions (from 3 mice) |
| | | | | Matrix vs Patch | Two-tailed unpaired t-test | 38 | t = 7.21 | **< 0.0001** | |
| S2B | Gαolf - Pyk2$^{+/+}$ | 100 | 5.23 | | | | | | 6 mice |
| | Gαolf - Pyk2$^{-/-}$ | 100.63 | 3.18 | | | | | | 6 mice |
| | | | | Gαolf - Pyk2$^{+/+}$ vs Pyk2$^{-/-}$ | Two-tailed unpaired t-test | 10 | t = 0.10 | 0.9206 | |
| | DARPP-32 - Pyk2$^{+/+}$ | 100 | 5.19 | | | | | | 6 mice |
| | DARPP-32 - Pyk2$^{-/-}$ | 106.28 | 3.52 | | | | | | 6 mice |
| | | | | DARPP-32 - Pyk2$^{+/+}$ vs Pyk2$^{-/-}$ | Two-tailed unpaired t-test | 10 | t = 1.00 | 0.3405 | |

| | | | | | | | | | |
|---|---|---|---|---|---|---|---|---|---|
| S2D | Synapsin - Pyk2$^{+/+}$ | 100 | 2.79 | | | | | | 6 mice |
| | Synapsin - Pyk2$^{-/-}$ | 94.57 | 3.99 | | | | | | 8 mice |
| | | | | Synapsin - Pyk2$^{+/+}$ vs Pyk2$^{-/-}$ | Two-tailed unpaired t-test | 12 | t = 1.04 | 0.3193 | |
| | TH - Pyk2$^{+/+}$ | 100 | 6.00 | | | | | | 6 mice |
| | TH - Pyk2$^{-/-}$ | 92.00 | 5.39 | | | | | | 8 mice |
| | | | | TH - Pyk2$^{+/+}$ vs Pyk2$^{-/-}$ | Two-tailed unpaired t-test | 10 | t = 0.99 | 0.3449 | |
| S3A | Pyk2$^{+/+}$ | | | | | | | | 9 mice |
| | Pyk2$^{-/-}$ | | | | | | | | 9 mice |
| | | | | | Two-way ANOVA:<br>Interaction<br>Trial<br>Genotype | <br>11<br>11<br>1 | <br>F = 0.39<br>F = 14.03<br>F = 2.57 | <br>0.9576<br>**< 0.0001**<br>0.1108 | |
| S3B | Pyk2$^{f/f;NAc,GFP}$ | | | | | | | | 17 mice |
| | Pyk2$^{f/f;NAc,GFP-Cre}$ | | | | | | | | 14 mice |
| | | | | | Two-way ANOVA:<br>Interaction<br>Trial<br>Genotype | <br>11<br>11<br>1 | <br>F = 0.44<br>F = 12.20<br>F = 1.83 | <br>0.9377<br>**< 0.0001**<br>0.1775 | |
| S3C | Pyk2$^{f/f;DS,GFP}$ | | | | | | | | 8 mice |
| | Pyk2$^{f/f;DS,GFP-Cre}$ | | | | | | | | 7 mice |
| | | | | | Two-way ANOVA:<br>Interaction<br>Trial<br>Genotype | <br>11<br>11<br>1 | <br>F = 0.70<br>F = 4.99<br>F = 1.07 | <br>0.7394<br>**< 0.0001**<br>0.3034 | |
| S3D | Pyk2$^{f/f}$ | | | | | | | | 14 mice |
| | Pyk2$^{f/f;D1::Cre}$ | | | | | | | | 13 mice |
| | | | | | Two-way ANOVA:<br>Interaction<br>Trial<br>Genotype | <br>11<br>11<br>1 | <br>F = 0.59<br>F = 22.46<br>F = 0.02 | <br>0.836<br>**< 0.0001**<br>0.8827 | |
| S3E | Pyk2$^{f/f}$ | | | | | | | | 14 mice |
| | Pyk2$^{f/f;A2A::Cre}$ | | | | | | | | 12 mice |
| | | | | | Two-way ANOVA:<br>Interaction<br>Trial<br>Genotype | <br>11<br>11<br>1 | <br>F = 0.27<br>F = 14.47<br>F = 0.84 | <br>0.991<br>**< 0.0001**<br>0.3605 | |

| | | | | | | | | | |
|---|---|---|---|---|---|---|---|---|---|
| S4A | 1st injection - $Pyk2^{f/f;NAc,GFP}$ | 106.02 | 11.90 | | | | | | 11 mice |
| | 2nd injection - $Pyk2^{f/f;NAc,GFP}$ | 136.72 | 11.76 | | | | | | 10 mice |
| | 1st injection - $Pyk2^{f/f;NAc,GFP-Cre}$ | 60.59 | 10.64 | | | | | | 11 mice |
| | 2nd injection - $Pyk2^{f/f;NAc,GFP-Cre}$ | 107.09 | 12.42 | | | | | | 10 mice |
| | | | | | Two-way ANOVA: | | | | |
| | | | | | Interaction | 1 | F = 0.84 | 0.3712 | |
| | | | | | AAV type | 1 | F = 6.99 | **0.016** | |
| | | | | | Injection | 1 | F = 20.05 | **0.0003** | |
| | | | | | Subjects (matching) | 19 | F = 2.71 | **0.0177** | |
| | | | | $Pyk2^{f/f;NAc,GFP}$: 1st inj vs 2nd inj | Sidak's multiple comparisons test | 19 | t = 2.58 | **0.0363** | |
| | | | | $Pyk2^{f/f;NAc,GFP-Cre}$: 1st inj vs 2nd inj | Sidak's multiple comparisons test | 19 | t = 3.73 | **0.0029** | |
| S4B | 1st injection - $Pyk2^{f/f;DS,GFP}$ | 78.99 | 10.00 | | | | | | 8 mice |
| | 2nd injection - $Pyk2^{f/f;DS,GFP}$ | 136.62 | 13.62 | | | | | | 7 mice |
| | 1st injection - $Pyk2^{f/f;DS,GFP-Cre}$ | 85.11 | 17.02 | | | | | | 8 mice |
| | 2nd injection - $Pyk2^{f/f;DS,GFP-Cre}$ | 140.06 | 9.79 | | | | | | 7 mice |
| | | | | | Two-way ANOVA: | | | | |
| | | | | | Interaction | 1 | F = 0.01 | 0.9044 | |
| | | | | | AAV type | 1 | F = 0.11 | 0.7478 | |
| | | | | | Injection | 1 | F = 26.55 | **0.0002** | |
| | | | | | Subjects (matching) | 13 | F = 1.78 | 0.1562 | |
| | | | | $Pyk2^{f/f;DS,GFP}$: 1st inj vs 2nd inj | Sidak's multiple comparisons test | 13 | t = 3.86 | **0.0039** | |
| | | | | $Pyk2^{f/f;DS,GFP-Cre}$: 1st inj vs 2nd inj | Sidak's multiple comparisons test | 13 | t = 3.44 | **0.0087** | |
| S5A | 1st injection - $Pyk2^{f/f}$ | 117.36 | 9.77 | | | | | | 25 mice |
| | 2nd injection - $Pyk2^{f/f}$ | 168.15 | 9.16 | | | | | | 23 mice |
| | 1st injection - $Pyk2^{f/f;D1::Cre}$ | 101.92 | 8.24 | | | | | | 25 mice |
| | 2nd injection - $Pyk2^{f/f;D1::Cre}$ | 132.43 | 6.76 | | | | | | 23 mice |
| | | | | | Two-way ANOVA: | | | | |
| | | | | | Interaction | 1 | F = 2.65 | 0.1103 | |
| | | | | | Genotype | 1 | F = 5.89 | **0.0191** | |
| | | | | | Injection | 1 | F = 42.60 | **< 0.0001** | |
| | | | | | Subjects (matching) | 46 | F = 2.86 | **0.0003** | |
| | | | | $Pyk2^{f/f}$: 1st inj vs 2nd inj | Sidak's multiple comparisons test | 46 | t = 5.89 | **< 0.0001** | |
| | | | | $Pyk2^{f/f;D1::Cre}$: 1st inj vs 2nd inj | Sidak's multiple comparisons test | 46 | t = 3.39 | **0.0029** | |
| S5B | 1st injection - $Pyk2^{f/f}$ | 94.73 | 10.02 | | | | | | 14 mice |
| | 2nd injection - $Pyk2^{f/f}$ | 145.63 | 11.84 | | | | | | 12 mice |
| | 1st injection - $Pyk2^{f/f;A2A::Cre}$ | 97.61 | 16.40 | | | | | | 14 mice |
| | 2nd injection - $Pyk2^{f/f;A2A::Cre}$ | 138.78 | 13.16 | | | | | | 12 mice |
| | | | | | Two-way ANOVA: | | | | |
| | | | | | Interaction | 1 | F = 0.43 | 0.5172 | |
| | | | | | Genotype | 1 | F = 0.01 | 0.9057 | |
| | | | | | Injection | 1 | F = 38.68 | **< 0.0001** | |
| | | | | | Subjects (matching) | 24 | F = 5.01 | **< 0.0001** | |
| | | | | $Pyk2^{+/+}$: 1st inj vs 2nd inj | Sidak's multiple comparisons test | 24 | t = 5.06 | **< 0.0001** | |
| | | | | $Pyk2^{f/f;A2A::Cre}$: 1st inj vs 2nd inj | Sidak's multiple comparisons test | 24 | t = 3.79 | **0.0018** | |

| | | | | | | | | | |
|---|---|---|---|---|---|---|---|---|---|
| S6B left | sal - Pyk2$^{f/f}$ | 100.00 | 7.23 | | | | | | 10 mice |
| | coc - Pyk2$^{f/f}$ | 123.55 | 3.85 | | | | | | 11 mice |
| | sal - Pyk2$^{f/f;D1::Cre}$ | 103.95 | 2.78 | | | | | | 13 mice |
| | coc - Pyk2$^{f/f;D1::Cre}$ | 115.20 | 6.61 | | | | | | 12 mice |
| | | | | | Two-way ANOVA: | | | | |
| | | | | | Interaction | 1 | F = 1.36 | 0.2498 | |
| | | | | | Genotype | 1 | F = 0.17 | 0.6789 | |
| | | | | | Drug | 1 | F = 10.89 | **0.0020** | |
| | | | | Pyk2$^{f/f}$: sal vs coc | Sidak's multiple comparisons test | 42 | t = 3.03 | **0.0084** | |
| | | | | Pyk2$^{f/f;D1::Cre}$: sal vs coc | Sidak's multiple comparisons test | 42 | t = 1.58 | 0.2291 | |
| S6B right | sal - Pyk2$^{f/f}$ | 100.00 | 6.12 | | | | | | 10 mice |
| | coc - Pyk2$^{f/f}$ | 105.97 | 4.89 | | | | | | 11 mice |
| | sal - Pyk2$^{f/f;D1::Cre}$ | 99.63 | 3.42 | | | | | | 12 mice |
| | coc - Pyk2$^{f/f;D1::Cre}$ | 100.99 | 6.36 | | | | | | 12 mice |
| | | | | | Two-way ANOVA: | | | | |
| | | | | | Interaction | 1 | F = 0.19 | 0.6662 | |
| | | | | | Genotype | 1 | F = 0.25 | 0.6165 | |
| | | | | | Drug | 1 | F = 0.48 | 0.4933 | |
| | | | | Pyk2$^{f/f}$: sal vs coc | Sidak's multiple comparisons test | 41 | t = 0.77 | 0.6925 | |
| | | | | Pyk2$^{f/f;D1::Cre}$: sal vs coc | Sidak's multiple comparisons test | 41 | t = 0.19 | 0.978 | |
| S6C left | sal - Pyk2$^{f/f}$ | 100.00 | 3.82 | | | | | | 5 mice |
| | coc - Pyk2$^{f/f}$ | 131.15 | 9.70 | | | | | | 6 mice |
| | sal - Pyk2$^{f/f;D1::Cre}$ | 108.69 | 7.61 | | | | | | 6 mice |
| | coc - Pyk2$^{f/f;D1::Cre}$ | 135.62 | 7.83 | | | | | | 6 mice |
| | | | | | Two-way ANOVA: | | | | |
| | | | | | Interaction | 1 | F = 0.07 | 0.7905 | |
| | | | | | Genotype | 1 | F = 0.70 | 0.4124 | |
| | | | | | Drug | 1 | F = 13.69 | **0.0015** | |
| | | | | Pyk2$^{f/f}$: sal vs coc | Sidak's multiple comparisons test | 19 | t = 2.74 | **0.0257** | |
| | | | | Pyk2$^{f/f;D1::Cre}$: sal vs coc | Sidak's multiple comparisons test | 19 | t = 2.49 | **0.0443** | |
| S6C right | sal - Pyk2$^{f/f}$ | 100.00 | 4.02 | | | | | | 5 mice |
| | coc - Pyk2$^{f/f}$ | 102.00 | 4.58 | | | | | | 6 mice |
| | sal - Pyk2$^{f/f;D1::Cre}$ | 100.37 | 5.43 | | | | | | 6 mice |
| | coc - Pyk2$^{f/f;D1::Cre}$ | 99.56 | 7.15 | | | | | | 6 mice |
| | | | | | Two-way ANOVA: | | | | |
| | | | | | Interaction | 1 | F = 0.06 | 0.8034 | |
| | | | | | Genotype | 1 | F = 0.03 | 0.8541 | |
| | | | | | Drug | 1 | F = 0.01 | 0.9161 | |
| | | | | Pyk2$^{f/f}$: sal vs coc | Sidak's multiple comparisons test | 19 | t = 0.25 | 0.9626 | |
| | | | | Pyk2$^{f/f;D1::Cre}$: sal vs coc | Sidak's multiple comparisons test | 19 | t = 0.11 | 0.9931 | |

| | | | | | | | | | |
|---|---|---|---|---|---|---|---|---|---|
| S7A | Day 0 - Pyk2$^{+/+}$ | 402.42 | 26.47 | | | | | | 10 mice |
| | Day 7 - Pyk2$^{+/+}$ | 556.80 | 28.46 | | | | | | 9 mice |
| | Day 0 - Pyk2$^{-/-}$ | 374.17 | 30.01 | | | | | | 10 mice |
| | Day 7 - Pyk2$^{-/-}$ | 515.37 | 36.99 | | | | | | 9 mice |
| | | | | | Two-way ANOVA: | | | | |
| | | | | | Interaction | 1 | F = 0.04 | 0.8526 | |
| | | | | | Genotype | 1 | F = 1.90 | 0.1859 | |
| | | | | | Injection | 1 | F = 17.90 | **0.0006** | |
| | | | | | Subjects (matching) | 17 | F = 0.52 | 0.9038 | |
| | | | | Pyk2$^{+/+}$: Day 0 vs Day 7 | Sidak's multiple comparisons test | 17 | t = 3.21 | **0.0102** | |
| | | | | Pyk2$^{-/-}$: Day 0 vs Day 7 | Sidak's multiple comparisons test | 17 | t = 2.79 | **0.0252** | |
| S7B | Pyk2$^{+/+}$ | 154.4 | 46.39 | | | | | | 10 mice |
| | Pyk2$^{-/-}$ | 141.2 | 52.61 | | | | | | 9 mice |
| | | | | Pyk2$^{+/+}$ vs Pyk2$^{-/-}$ | Two-tailed unpaired t-test | 17 | t = 0.19 | 0.8526 | |
| S7C | Pyk2$^{f/f;NAc,GFP}$ | 114.4 | 30.14 | | | | | | 18 mice |
| | Pyk2$^{f/f;NAc,GFP\text{-}Cre}$ | 83.26 | 32.02 | | | | | | 15 mice |
| | | | | Pyk2$^{f/f;NAc,GFP}$ vs Pyk2$^{f/f;NAc,GFP\text{-}Cre}$ | Two-tailed unpaired t-test | 31 | t = 0.71 | 0.4855 | |
| S7D | Pyk2$^{f/f;DS,GFP}$ | 142.4 | 46.55 | | | | | | 8 mice |
| | Pyk2$^{f/f;DS,GFP\text{-}Cre}$ | 145.8 | 41.61 | | | | | | 7 mice |
| | | | | Pyk2$^{f/f;DS,GFP}$ vs Pyk2$^{f/f;DS,GFP\text{-}Cre}$ | Two-tailed unpaired t-test | 13 | t = 0.05 | 0.958 | |
| S7E | Pyk2$^{f/f}$ | 135.4 | 39.96 | | | | | | 13 mice |
| | Pyk2$^{f/f;D1::Cre}$ | 160.4 | 21.3 | | | | | | 12 mice |
| | | | | Pyk2$^{+/+}$ vs Pyk2$^{f/f;D1::Cre}$ | Two-tailed unpaired t-test | 23 | t = 0.54 | 0.5948 | |
| S7F | Pyk2$^{f/f}$ | 170 | 26.14 | | | | | | 14 mice |
| | Pyk2$^{f/f;A2A::Cre}$ | 136.5 | 34.62 | | | | | | 12 mice |
| | | | | Pyk2$^{+/+}$ vs Pyk2$^{f/f;A2A::Cre}$ | Two-tailed unpaired t-test | 24 | t = 0.78 | 0.4405 | |
| S8A | Pyk2$^{f/f}$ | | | | | | | | 13 mice |
| | Pyk2$^{f/f;D1::Cre}$ | | | | | | | | 12 mice |
| | | | | | Two-way ANOVA: | | | | |
| | | | | | Interaction | 17 | F = 0.95 | 0.51 | |
| | | | | | Time | 17 | F = 19.11 | **< 0.0001** | |
| | | | | | Genotype | 1 | F = 0.00 | 0.9499 | |
| S8B | Pyk2$^{f/f}$ | | | | | | | | 12 mice |
| | Pyk2$^{f/f;D1::Cre}$ | | | | | | | | 12 mice |
| | | | | | Two-way ANOVA: | | | | |
| | | | | | Interaction | 17 | F = 0.60 | 0.8922 | |
| | | | | | Time | 17 | F = 20.46 | **< 0.0001** | |
| | | | | | Genotype | 1 | F = 0.44 | 0.5052 | |
| S8C | Pyk2$^{f/f}$ | 1.11 | 0.04 | | | | | | 12 mice |
| | Pyk2$^{f/f;D1::Cre}$ | 1.16 | 0.05 | | | | | | 12 mice |
| | | | | Pyk2$^{+/+}$ vs Pyk2$^{f/f;D1::Cre}$ | Two-tailed unpaired t-test | 22 | t = 0.83 | 0.413 | |
