## Supplementary figures for "Pyk2 in D1 receptor-expressing neurons of the nucleus accumbens modulates the acute locomotor effects of cocaine"

### Supplementary Figures
# S1-S8

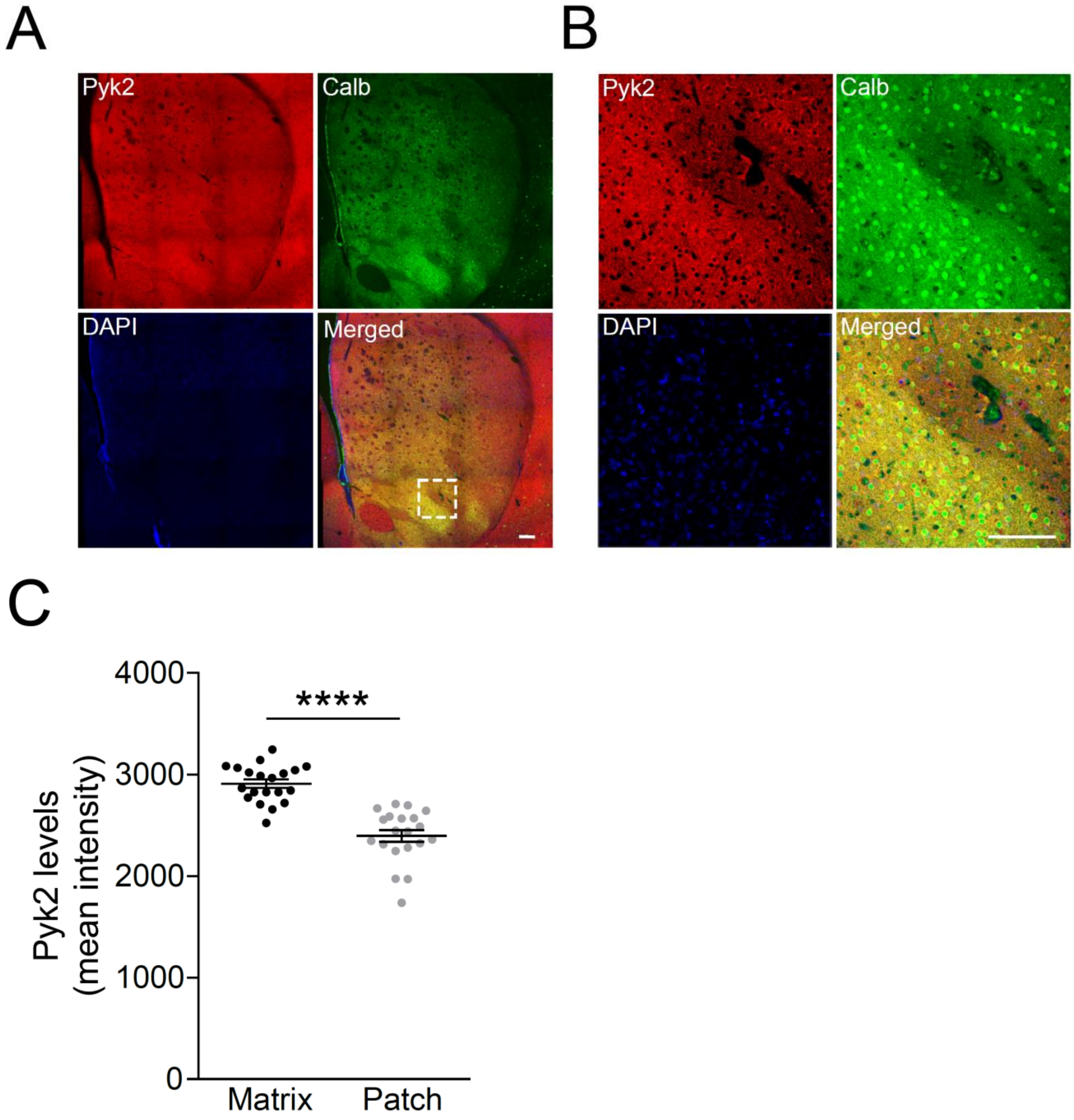


**Figure S1: Enrichment of Pyk2 in the striatal matrix. a**, Distribution of Pyk2 (red) and calbindin (green) immunoreactivity, and DAPI (blue) in the striatum of wild-type mice. Scale bar: 150 µm. **b**, Zoom in the white square indicated in **a. c**, Pyk2 labelling intensity in striatal matrix as defined as calbindin-enriched areas. Values are means ± SEM. Unpaired t-test, ****p<0.0001. For number of replicates and statistical analysis, see **Supplementary Table 2**.

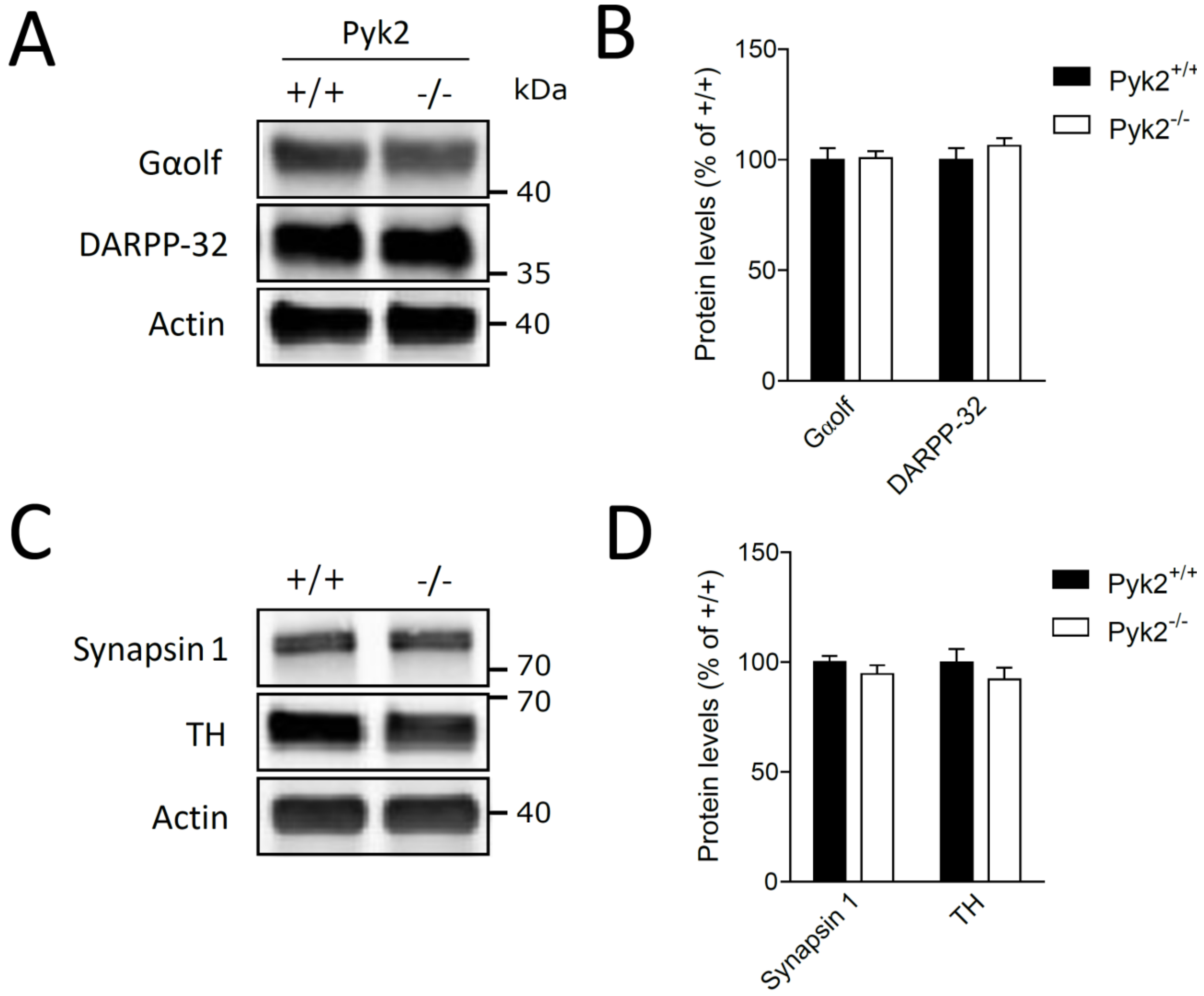


**Figure S2: Striatal proteins levels in Pyk2-deficient mice. a**, Immunoblotting analysis of Gαolf, DARPP-32, and actin as loading control in 4-month-old Pyk2$^{+/+}$ and Pyk2$^{-/-}$ mice. **b**, Densitometry quantification of results as in **a**. Data were normalized to actin for each sample and expressed as percentage of wild type averge. **c**, Synapsin 1, tyrosine hydroxylase, and actin were analyzed by immunoblotting. **d**, Results as in **c** were quantified and analyzed as indicated in **b.** In **d** and **d**, statistical analysis was done with unpaired t-test. No significant difference. In all graphs, values are means + SEM. For number of mice and detailed statistical analysis, see S**upplementary Table 2**.

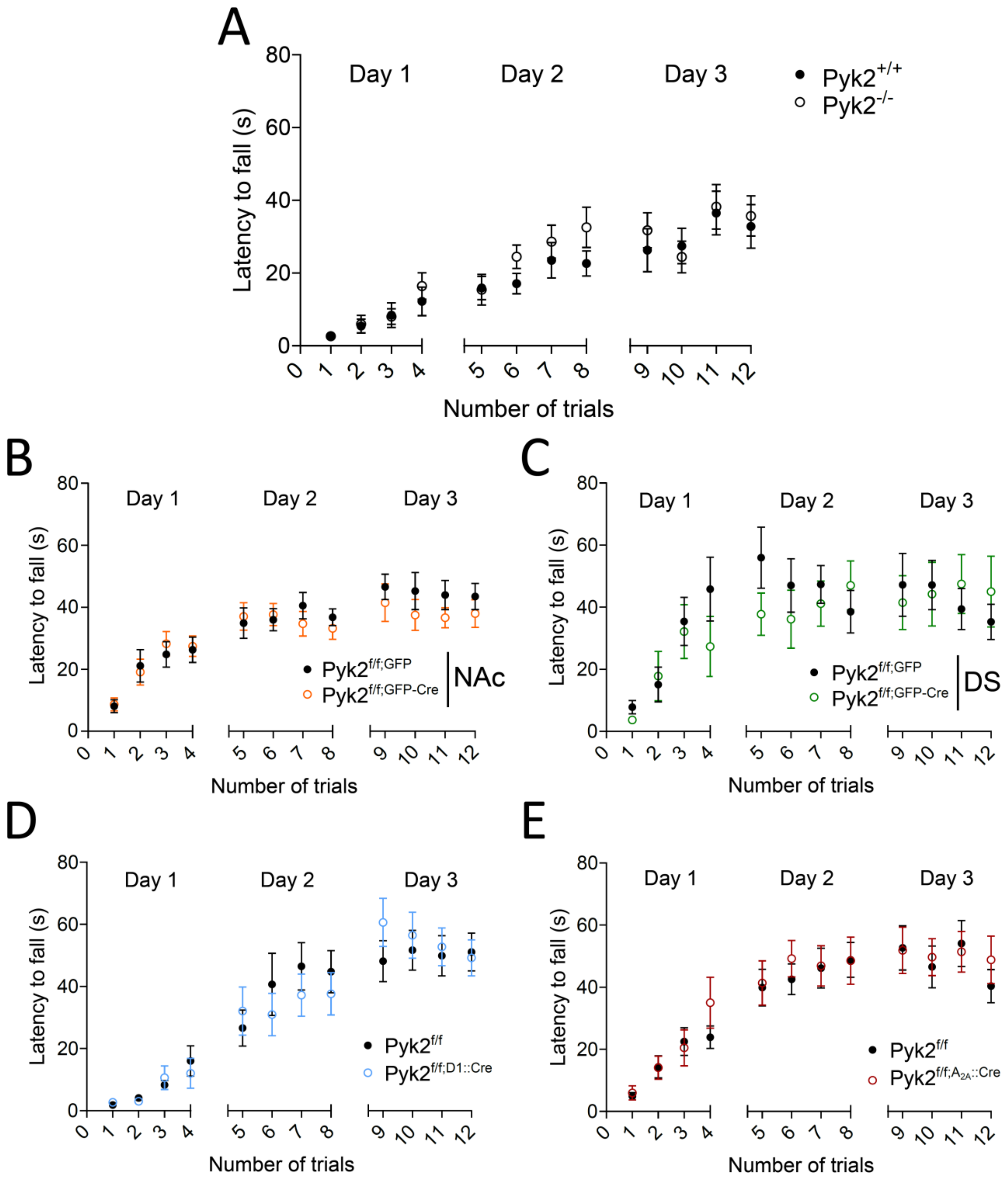


**Figure S3: Motor skill learning on the accelerated rotarod is not altered in Pyk2 mutant mice.** Latency to fall on the accelerating rotarod for each trial at days 1, 2, and 3 for the various Pyk2 mutant mice and their respective matched controls. **a**, Pyk2$^{+/+}$ and Pyk2$^{-/-}$. **b**, Pyk2$^{f/f;NAc,GFP}$ and Pyk2$^{f/f;NAc,GFP\text{-}Cre}$. **c**, Pyk2$^{f/f;DS,GFP}$ and Pyk2$^{f/f;DS,GPF\text{-}Cre}$. **d**, Pyk2$^{f/f}$ and Pyk2$^{f/f;D1::Cre}$. **e**, Pyk2$^{f/f}$ and Pyk2$^{f/f;A2A::Cre}$ mice. Values are means ± SEM. Two-way ANOVA, genotype or AAV effect, no difference. For number of mice and detailed statistical analysis, see **Supplementary Table 1**.

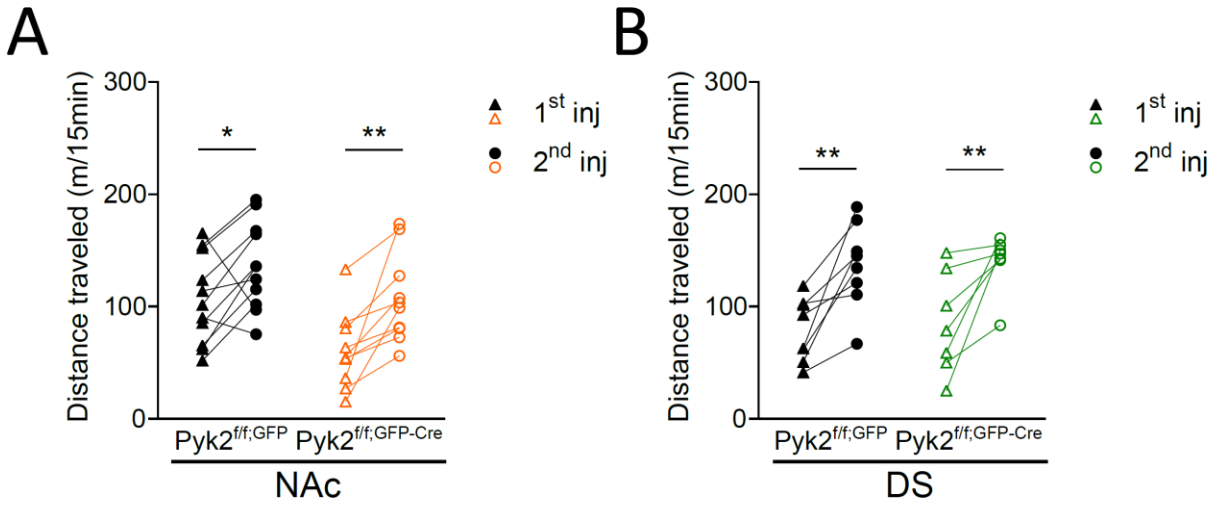


**Figure S4: Pyk2 deletion in the NAc or in the DS does not alter locomotor sensitization.** Thirty minutes after being placed in a circular open space, mice received an intraperitoneal injection of 20 mg/kg cocaine on Day 1 ($1^{st}$ injection) and Day 14 ($2^{nd}$ injection), as described in the Materials and Methods. The distances traveled during the 15 first minutes following the injection is plotted for each mouse (see average locomotor activity in **Fig. 3**). **a**, Pyk2$^{f/f;NAc,GFP}$ and Pyk2$^{f/f;NAc,GFP\text{-}Cre}$ mice. **b**, Pyk2$^{f/f;DS,GFP}$ and Pyk2$^{f/f;DS,GPF\text{-}Cre}$ mice. In **a** and **b**, statistical analysis with two-way ANOVA and Sidak's multiple comparisons post hoc test, **$p < 0.01$, *$p < 0.05$. For number of mice and detailed statistical analysis, see **Supplementary Table 1**.

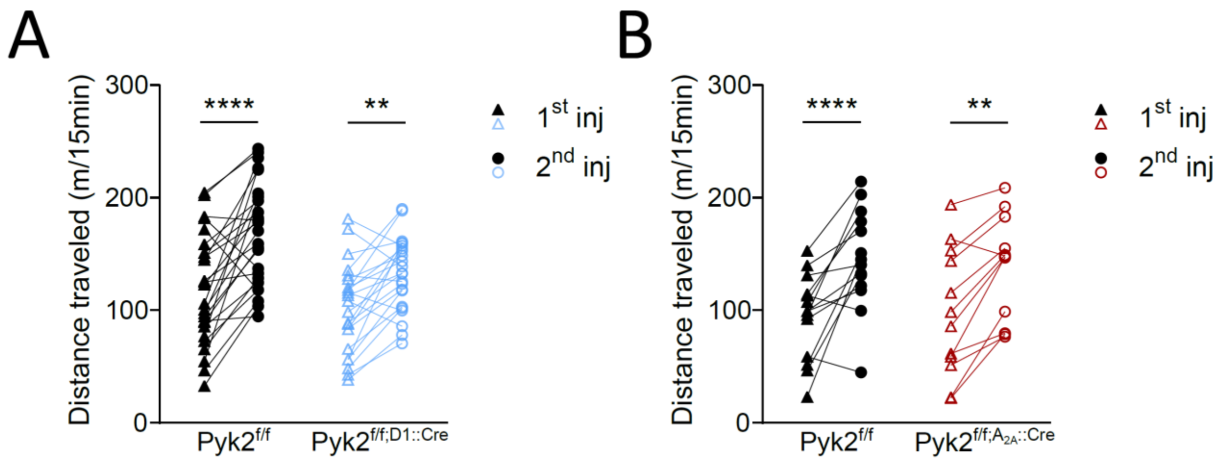


**Figure S5: Pyk2 deletion in D1 or in $A_{2A}$ neurons does not alter locomotor sensitization.** Thirty minutes after being placed in a circular open space, mice received an intraperitoneal injection of 20 mg/kg cocaine on Day 1 ($1^{st}$ injection) and Day 14 ($2^{nd}$ injection), as described in the Materials and Methods. The distances traveled during the 15 first minutes following the injection is plotted for each mouse (see average locomotor activity in **Fig. 4**). **a**, Pyk2$^{f/f}$ and Pyk2$^{f/f;D1::Cre}$ mice. **b**, Pyk2$^{f/f}$ and Pyk2$^{f/f;A2A::Cre}$ mice. In **a** and **b**, statistical analysis with two-way ANOVA and Sidak's multiple comparisons post hoc test, ****$p < 0.0001$, **$p < 0.01$, *$p < 0.05$. For number of mice and detailed statistical analysis, see **Supplementary Table 1**.

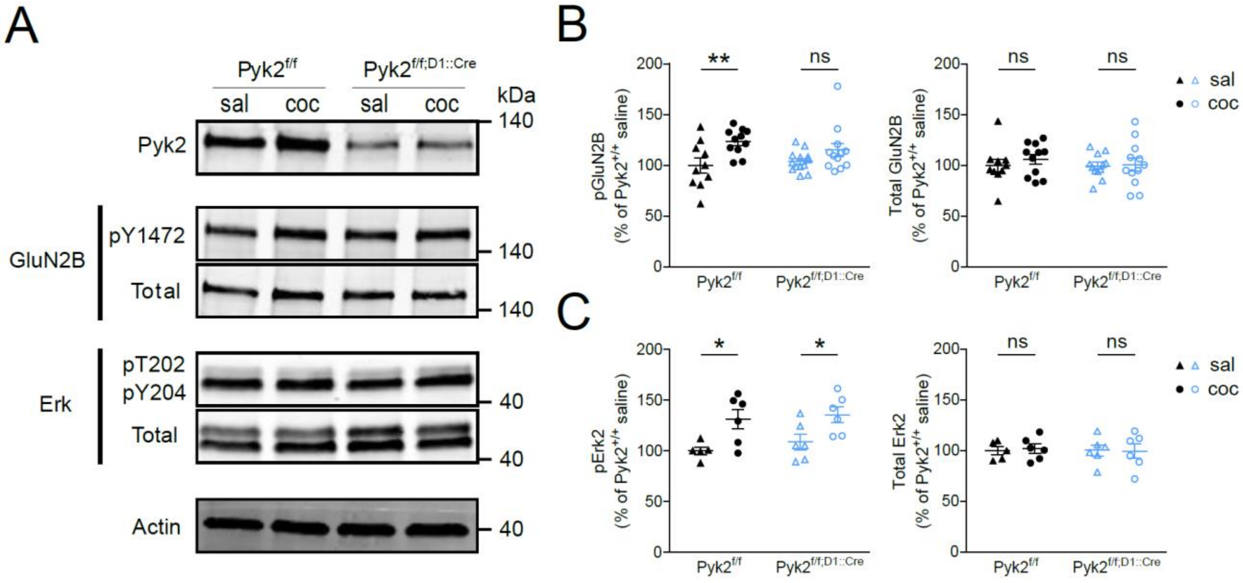


**Figure S6: Specific deletion of Pyk2 in D1 neurons decreases cocaine effects on GluN2B phosphorylation but not ERK. a** Immunoblotting analysis of Pyk2, GluN2B phosphorylated an Tyr-1472 (**pY1472**) and total levels (**Total**), ERK phosphorylated at Thr-202 and Tyr-204 (**pT202/pY204**) and total levels (**Total**), and actin as loading control in 3-month Pyk2$^{f/f}$ and Pyk2$^{f/f;D1::Cre}$ mice, 10 min after saline (**sal**) or cocaine (20 mg/kg, **coc**) i.p. injection . **b** Densitometry quantification of phospho (**left**) and total (**right**) GluN2B levels. **c** Densitometry quantification of phosphorylated (**left**) and total (**right**) ERK levels. In **b** and **c**, data were normalized to actin for each sample and expressed as percentages of the average in saline-injected Pyk2$^{f/f}$ mice. Statistical analysis was done with two-way ANOVA and Sidak's multiple comparisons post hoc test, *p < 0.05, *p < 0.05, not significant, ns. In all graphs, means +/- SEM are indicated. For number of mice and detailed statistical analysis, see **Supplementary Table 2**.

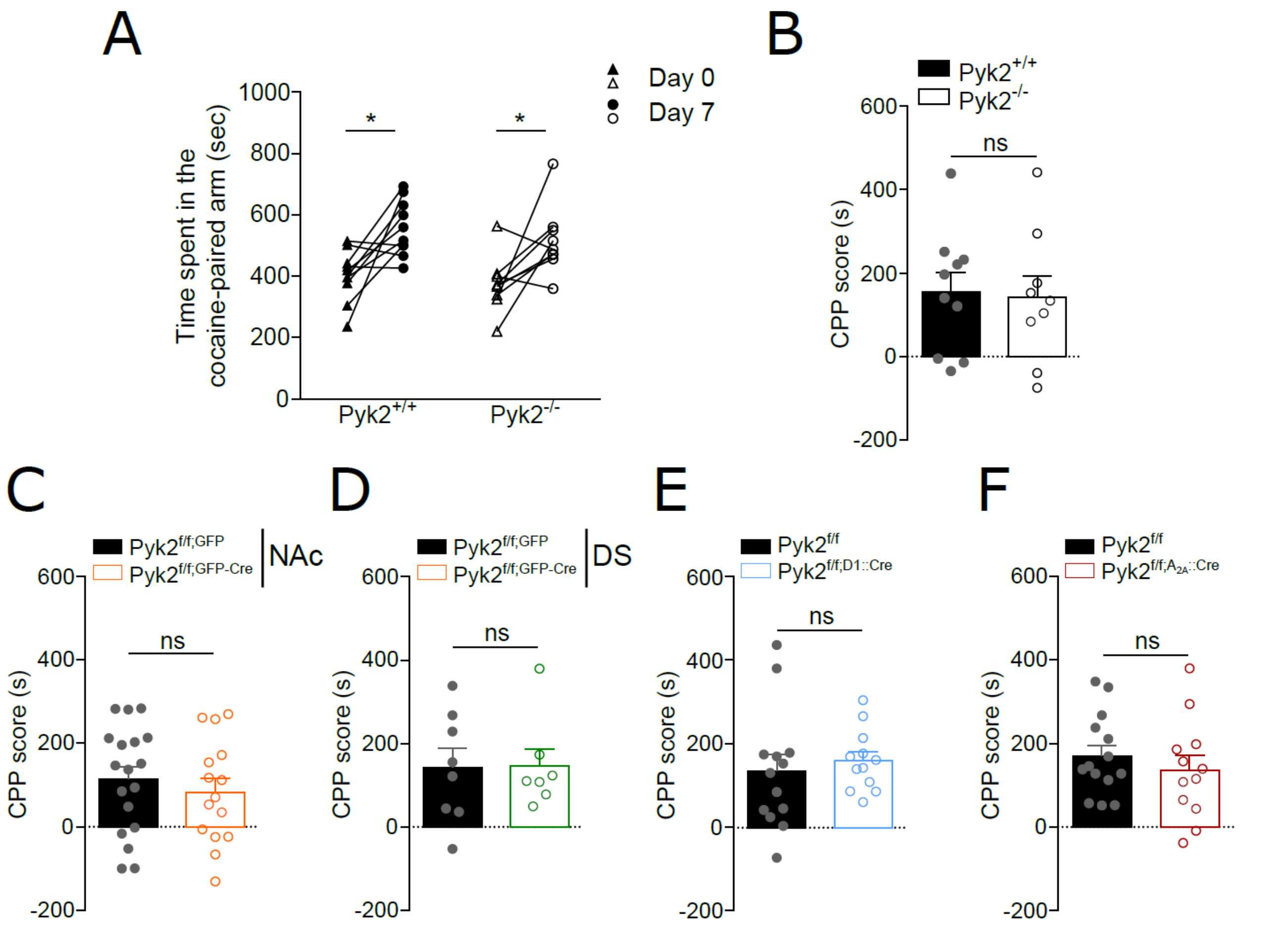


**Figure S7: Pyk2 deletion does not alter cocaine-conditioned place preference.** For each group of Pyk2 mutant mice and their respective matched controls, the time spent in the cocaine-paired arm before (Day 0) and after (Day 7) cocaine (15 mg/kg) conditioning was measured (see **Materials and Methods**). **a**, Time spent in the cocaine-paired arm in Pyk2$^{+/+}$ and Pyk2$^{-/-}$ mice. Two-way ANOVA and Sidak's multiple comparisons post hoc tests, *$p < 0.05$. **b**, The CPP score was calculated for each Pyk2$^{+/+}$ and Pyk2$^{-/-}$ mouse in **a** as the excess time spent in the cocaine-paired arm. **c-f**, Same CPP score as in B for Pyk2$^{f/f;NAc,GFP}$ and Pyk2$^{f/f;NAc,GFP-Cre}$ mice (**c**), Pyk2$^{f/f;DS,GFP}$ and Pyk2$^{f/f;DS,GPF-Cre}$, mice (**d**), Pyk2$^{f/f}$ and Pyk2$^{f/f;D1::Cre}$ (**e**), and Pyk2$^{f/f}$ and Pyk2$^{f/f;A2A::Cre}$ mice (**f**). **b-f**, Bars indicate means + SEM, unpaired t-test, ns, not significant. For number of mice and detailed statistical analysis, see **Supplementary Table 2**.

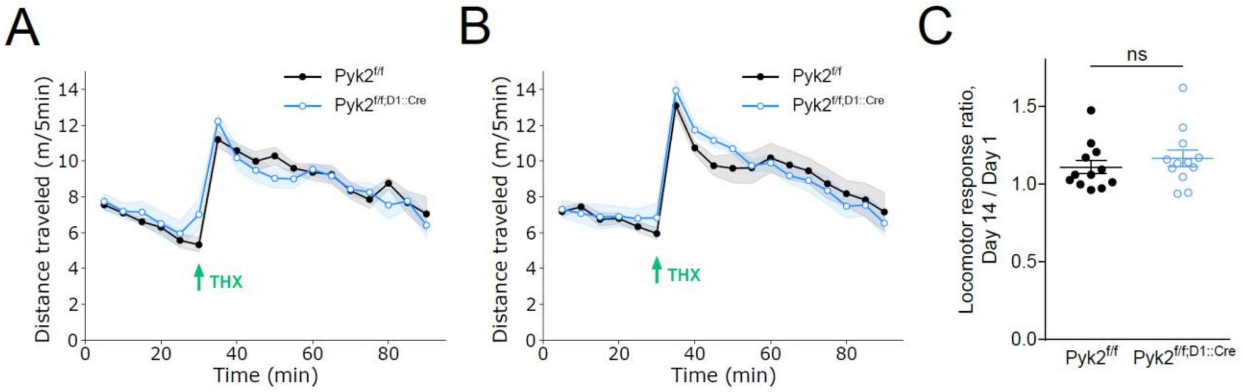


**Figure S8: Specific deletion of Pyk2 in D1 neurons does not alter the locomotor responses to trihexyphenidine. a-b**, Locomotor activity of Pyk2$^{f/f}$ and Pyk2$^{f/f;D1::Cre}$ mice after a first (**a**) and, 13 days later, a second (**b**) injection of trihexyphenidine (THX). THX (15 mg/kg i.p., arrow) was injected 30 min after mice were placed in the open field. Two-way ANOVA, no genotype effect. **c**, Sensitization during the first 10 min after SKF injection (data from **a** and **b**). **c**, Ratio of the distances traveled 0-15 min after the second (Day 14) / the first (Day 1) THX injections. Unpaired t-test, not significant, ns. In all graphs, means ± SEM, are indicated (shaded area in **a** and **b**). For detailed statistical analysis, see **Supplementary Table 2**.
